# The biodistribution and effect of post-exposure neutralising monoclonal antibody treatment in a mouse model of SARS-CoV-2 infection with viral spread to the brain

**DOI:** 10.64898/2026.05.27.728081

**Authors:** Janine Anja Schläpfer, Simon De Neck, Rebekah Penrice-Randal, Parul Sharma, Adam Kirby, Lee Tatham, Eduardo Gallardo-Toledo, Joanne Herriott, Edyta Kijak, Joanne Sharp, James P. Stewart, Andrew Owen, Anja Kipar

## Abstract

Ronapreve, a combination of two neutralising monoclonal antibodies, casirivimab and imdevimab, was amongst the authorised treatments against SARS-CoV-2 early in the COVID-19 pandemic. Ronapreve has lost some of its efficiency with the rise of new virus variants, however, it remains a valuable tool for experimental studies to gain insights into the mechanisms and effects of anti-viral drugs. In this study we combined morphological, pharmacokinetic and molecular approaches (including multiomics) to investigate the biodistribution of Ronapreve in the K18-hACE2 murine model of SARS-CoV-2 neuroinvasion, as well as possible consequences for the brain. We also investigated the effect of the treatment on the infection status. Our results showed that after intraperitoneal injection, Ronapreve accumulates in the serum and is unable to cross the blood-brain barrier, thus not reaching the brain parenchyma; treatment has only a minimal effect on the brain transcriptome, with no significant changes in the brain lipidome or metabolome. Nonetheless, post-exposure Ronapreve treatment resulted in reduced viral loads in the lung and, in particular, the brain, with markedly reduced tissue response in the brain, as shown by the transcriptomic analysis. The results suggest a peripheral mode of action of Ronapreve to block brain infection, possibly by lowering viral replication in the nasal epithelium, reducing a subsequent spread to the brain.

## Introduction

Neutralising monoclonal antibodies (mAbs) were amongst the earliest targeted antiviral therapeutics deployed during the Severe Acute Respiratory Syndrome Coronavirus 2 (SARS-CoV-2) pandemic, where they demonstrated efficacy in pre (*1*)- and post-exposure prophylaxis (*2*), early treatment (*3*), and in the treatment of hospitalised patients with severe coronavirus disease 2019 (COVID-19) (*4, 5*). However, with the successive emergence of Omicron variants and sub-variants, all mAbs lost varying degrees of neutralising capacity (*6–8*). This led to the withdrawal of their regulatory authorisations and the discontinuation of clinical use (*9–12*). Despite the lack of clinical utility against post-Omicron SARS-CoV-2 variants, mAbs remain valuable tools for understanding the pharmacokinetics (PK), biodistribution, and antiviral behaviour of large-molecule biologics in acute respiratory viral infections. In the K18-hACE2 mouse model, intranasal challenge with pathogenic SARS-CoV-2 variants (e.g., ancestral, Alpha, Beta, Delta isolates) results in neuroinvasion with high frequency (*13–17*). The virus mainly spreads from the olfactory epithelium to the olfactory bulb, likely via axonal transport, and subsequently disseminates widely in the brain (*13, 15, 18*). Notably, in this model, the Omicron (BA.1) variant lacked this neuroinvasive phenotype (*13, 19–22*). The potential for such viral neuroinvasion and associated neuroinflammation is of significant interest, as persistent neurological symptoms, including cognitive impairment (‘brain fog’) and chronic fatigue, are hallmarks of Long COVID, highlighting the potential benefit of blocking virus CNS entry (*21, 23*). This pattern of olfactory-driven CNS entry is not unique to SARS-CoV-2 and is known for several other viruses, including influenza A, rabies and Japanese encephalitis virus (*24*). Multiple respiratory viruses, including influenza A (*24*), respiratory syncytial virus (*25*) and Nipah virus (*26*), use the olfactory mucosa as a mechanism for neuroinvasion in animal models. These parallels underscore the broader relevance of understanding how antiviral antibody exposure contributes to blocking neuroinvasion.

Previously, we demonstrated that intraperitoneal administration of Ronapreve (400 µg), 24 hours post infection (hpi), substantially reduced pulmonary viral loads and prevented dissemination of the SARS-CoV-2 Delta variant to the brain in a K18-hACE2 mouse model, suggesting a marked neuroprotective effect (*20*). Notably, we also observed an altered inflammatory response in the lungs following Ronapreve treatment, characterised by a granulomatous reaction likely resulting from focal recruitment of macrophages into the parenchyma in response to antibody-opsonised virus, indicating that mAb-mediated opsonisation drives measurable tissue responses. However, the study did not clarify whether protection was mediated by direct antibody penetration across the blood-brain barrier (BBB) or through suppression of viral replication at early peripheral sites of infection. IgG antibodies exhibit very low permeability across the intact BBB, making it uncertain whether systemically administered neutralising mAbs can access the neuroparenchyma at effective concentrations (*27, 28*). These findings raised questions about the biodistribution of mAbs following systemic administration and their potential effects independent of or in addition to SARS-CoV-2 infection, particularly on the brain. This constraint also makes mAbs a valuable tool for investigating how systemic antiviral activity can prevent CNS infection, where direct antibody exposure is expected to be limited.

Here, we conducted terminal PK analysis, tissue distribution profiling, and brain *in situ* immunostaining of casirivimab and imdevimab, the two mAbs comprising Ronapreve, at day 7 following infection with SARS-CoV-2 Delta in a model that reflects a treatment use case. These data enable assessment of whether Ronapreve reaches the brain parenchyma or whether neuroprotection arises from peripheral restriction of viral replication, particularly at the nasal mucosa, which likely serves as the critical gateway for CNS infection in this model.

## Material and Methods

### Virus

All animals in this study were infected with the SARS-CoV-2 Delta variant B.1.617.2 hCoV-19/England/SHEF-10E8F3B/2021 (GISAID accession number EPI_ISL_1731019), which was previously described (*20, 29*).

### Animal studies

All animal experiments were approved by the local University of Liverpool Animal Welfare and Ethical Review Body and carried out under UK Home Office Project Licences PP4715265. It was carried out in agreement with locally approved risk assessments and standard operating procedures.

This study was performed on 6-8-week-old female mice (16-24 g) carrying the human ACE2 gene controlled by keratin 18 promoter (K18-hACE2; formally B6.Cg-Tg(K18-ACE2)2Prlmn/J), purchased from Charles River Laboratories (Wilmington, MA, USA). Mice were housed in biosafety level 3 facilities under SPF barrier conditions in individually ventilated cages and a 12-hour light/dark cycle, at 21 °C ± 2 °C. Unrestricted access to food and water was ensured at all times. Mice were acclimatised for 7 days. They were randomly assigned to cohorts of 4-6 animals each. The experimental groups are listed in Supplemental Table S1.

#### Virus infection

For infection, mice were anesthetised with 3% isoflurane and inoculated intranasally with 50 µL of 10^3^ PFU of SARS-CoV-2 Delta (B.1.617.2) in phosphate buffered saline (PBS) on day 0. Control animals (mock infection) underwent the same procedure but received 50 µL PBS instead.

#### Ronapreve treatment

At 24 hpi (day 1), a cohort of virus infected (n=6) and mock infected (n=4) mice was treated with a single dose of 400 µg Ronapreve (casirivimab and imdevimab). Ronapreve was kindly provided by F. Hoffmann-La Roche Ltd (Switzerland). Prior to use, it was diluted in saline (100 µl) and administered via intraperitoneal (IP) injection, as previously described (*20*).

#### Untreated controls

One cohort of 5 virus infected mice and one cohort of 6 mock infected mice remained untreated and were not handled on Day 1; these served as controls.

All animals were weighed and clinically monitored daily throughout the experiments. At 7 days post infection (dpi), all mice were sacrificed via a lethal intraperitoneal injection of pentobarbitone, followed by cardiac puncture and immediate exsanguination from the heart. The blood was collected in serum separator tubes (Sarstedt, Germany). Animals were immediately dissected. Samples from the right lung and right frontal cortex were collected for virus RT-qPCR, and samples from the thalamus and the hypothalamus were collected for transcriptomic and metabolomic/lipidomic analyses, respectively. For the pharmacokinetic analysis, samples from the nasal turbinates, left frontal cortex, right lung, spleen, liver and kidney were collected. All these tissue samples were frozen at -80 °C until further processing. The left lung and the remaining brain (frontal cortex, brain stem with medulla oblongata, part of lobus occipitalis, cerebellum), liver, spleen and kidney were fixed in 10% buffered formalin for 48 h, then stored in 70% ethanol until processing for histological and immunohistological examination and RNA-*in situ* hybridisation (RNA-ISH).

### Determination of viral loads by RT-qPCR

Viral loads were quantified in samples from the right lung and the frontal cortex, using the GoTaq® Probe 1-Step RT-qPCR System (Promega). The nCOV_N1 primer/probe mix from the SARS-CoV-2 (2019-nCoV) CDC qPCR Probe Assay (IDT) and murine 18S primers were used to quantify SARS-COV-2, as previously described (*30*).

### Quantification of Ronapreve in mouse serum and tissues

Terminal cardiac bleeds from each animal were collected in serum separator tubes (Sarstedt, Germany), and serum was isolated following 1 h incubation of the blood at room temperature (RT). Isolated serum was mixed 1:4 with 0.5% v/v Triton X-100 in PBS and stored at −80 °C prior to analysis. For tissue sample preparation, tissue sections were rinsed in ice-cold PBS and transferred to Precellys CKmix lysing tubes (Bertin Technologies, France). Subsequently, 1 mL of 0.5% v/v Triton X-100 in PBS was added to each tissue sample. Homogenisation was performed using a bead mill homogeniser (Fisher Scientific, UK) at 3.5 m/s for up to two 30-second cycles. The resulting homogenate was centrifuged at 1000 x g for 15 min at 4 °C, and the supernatant was isolated and stored at -80 °C until analysis.

Casirivimab and imdevimab targets were quantified separately using ELISA (Abbexa, UK). Prior to the assay, both the serum and tissue supernatants were diluted with sterile dH_2_O. A standard curve was prepared by serial dilution ranging from 15.6 – 1000 ng/mL. All standards, blanks (standard diluent), and samples were prepared in duplicate and added to the 96-well ELISA microplates. The assays were subsequently conducted following the manufacturer’s instructions. Following incubation, optical density (OD) was measured using a BioTek Synergy microplate reader at 450 nm (Agilent, USA). An inverse correlation is observed between each mAb concentration and the measured OD. Average sample concentrations were calculated from OD values using the standard curves generated in Prism v.10.3 (GraphPad, USA). Tissue values were corrected for the initial sample weight.

### Histology, immunohistology and RNA-*in situ* hybridisation

The fixed left lung, brain, liver, kidney and spleen specimens were trimmed and routinely paraffin wax embedded. Consecutive sections (3-5 µm) were prepared and routinely stained with hematoxylin eosin (HE) for histological examination (all tissues), or subjected to immunohistological staining, using the horseradish peroxidase (HRP) method. For the detection of SARS-CoV antigen expression (sections from lung and brain), a rabbit anti-SARS-CoV nucleocapsid protein antibody (Rockland) was used, with incubation of the primary antibody at 4 °C overnight (diluted 1:6000 in dilution buffer), as previously described (*13, 29*). For the detection of casirivimab and imdevimab (sections from lung, brain, liver, kidney and spleen), sections were routinely deparaffinized and subjected to antigen retrieval in citrate buffer (pH 6.0) for 20 min at 98 °C, followed by overnight incubation at 4 °C with the primary antibodies (diluted to 10 µg/ml in dilution buffer, Agilent Dako): rabbit anti-casirivimab and rabbit anti-imdevimab (Abbexa, UK). This was followed by blocking of endogenous peroxidase (peroxidase block, Agilent Dako) for 10 min at RT and incubation with Envision+ rabbit (Agilent Dako) for 30 min at RT in an autostainer (Agilent Dako), with 3,3‘-diaminobenizidine (DAB) as chromogen. Sections were subsequently counterstained with hematoxylin. RNA *in situ* hybridisation (RNA-ISH) was performed on sections from lung and brain of the Ronapreve-treated mice using the RNAscope® ISH method (Advanced Cell Diagnostics (ACD), Newark, California) as previously described (*13*).

### Transcriptomic, metabolomic, and lipidomic analyses

Brain samples, from thalamus and hypothalamus, were collected and processed for bulk transcriptomic (thalamus), metabolomic (hypothalamus) and lipidomic (hypothalamus) analysis, respectively, as previously described (*14*) (Supplemental Figure S1). A list of the R packages used is available in the supplemental material (Supplemental Table S3A-C).

#### Transcriptomic

Extracted RNA was sequenced by Novogene (Novogene (UK) Company Limited, Cambridge, UK) using polyA enrichment and the NovaSeq X Plus (Illumina®, San Diego, California, USA). The sequencing reads were processed as previously described (*14*) and were imported in RStudio (v2026.01.0 Build 392) for downstream analysis with R (*31*). Using the edgeR package (v4.8.2 (*32*)), the genes were filtered and the library normalised, before performing differential expression analysis using the quasi-likelihood pipeline (glmQLFit() and glmQLFTest()); several comparison pairs were investigated using limma (v3.66.0) makeContrasts() function (*33*) (Supplemental Table S2A) and the results were visualised with volcano plots (EnhancedVolcano package, v1.28.2 (*34*)). Differential genes were defined as having at least a 4-fold difference (log2FC) compared to the reference cohort, and a false discovery rate (FDR) below 0.05. Enrichment analysis was carried out in clusterProfiler (v4.18.4 (*35–37*)) using the Gene Ontology (GO) database (*38, 39*), with either the Gene Set Enrichment Analysis (GSEA) method, or the Over-Representation Analysis method. Using PCAtools (v2.22.1 (*40*)), the transcriptomic profiles and clustering of the four cohorts were investigated, while the clustering of genes of interest was visualised with pheatmap (v1.0.13 (*41*)).

#### Metabolomic and lipidomic

The samples collection and processing and the downstream analysis in R have been previously described (*14*). Briefly, the metabolomic dataset was analysed with the TidyMass pipeline (v2.0.10 (*42*)), while the lipidomic dataset was analysed with the LipidSigR pipeline (v1.0.4 (*43*)). The various comparison pairs investigated are detailed in Supplemental Table S2B.

### Statistical analysis

#### Weight curves

The weight curves were obtained by plotting the variation in bodyweight for each cohort, with weight expressed as a percentage of the original bodyweight at day 0 (RStudio v2025.05.1 Build 513; R v4.5.1). Using the Rmisc::summarySE() function (v1.5.1) (*44*), at each timepoint the number of animals (count), the mean bodyweight, the standard deviation, the standard error of the mean, and the confidence interval were computed for each cohort. The mean bodyweight was plotted with ggplot2 (v4.0.0) (*45*), with error bars calculated as standard errors (ggplot2:: geom_errorbar()). A list of the packages used is available in Supplemental Table S4.

#### PCR for viral RNA in the brain and lung

Statistical analyses were undertaken to compare the PCR values between the Delta-infected and the Delta-infected and Ronapreve-treated cohorts, for both the lung and the brain. The statistical analyses were undertaken in RStudio (2025.05.1 Build 513) with R (version 4.5.1, 2025-06-13 ucrt) using the PCR values. The normality of the data for each cohort and organ was assessed using the Shapiro-Wilk test (rstatix::shapiro_test(), v0.7.2); all data were considered as normally distributed (Supplemental Table S5). The equality of the variance was assessed with Fisher’s F test (var.test); for each organ, the variance of both samples was not equal (Supplemental Table S5). As the variances are not equal and the sample sizes are small, the statistical analysis were done with a Mann-Whitney U test (ggpubr::compare_means(method = “wilcox.test), v0.6.1) (*46, 47*). The individual PCR values for each cohort and organ were plotted in the form of boxplot as raw data (ggpubr::ggboxplot(add = “jitter”), v0.6.1), and a log10 transformation was applied on the y axis to aid visualisation (ggplot2::scale_y_log10(), v4.0.0). A list of the packages used is available in Supplemental Table S4.

#### PK studies

The Shapiro-Wilk test was used to assess the normality of data distribution. Two-tailed Mann-Whitney tests were subsequently applied to compare groups. Statistical significance was set at p < 0.05. All statistical analyses were performed using GraphPad Prism v. 10.3.

## Results

### Body weights

Body weight was measured during the experiment as an indicator of health and is shown in Figure 1 in relation to the initial weight (Day 0; day of infection). Untreated infected mice showed the typical weight loss pattern observed with SARS-CoV-2 Delta infection (*30*), with the first drop in weight observed at 4 dpi. By the time of scheduled sacrifice at 7 dpi, the weight loss had partly reached the 20% mark, meeting the clinical endpoint.

**Figure 1.**
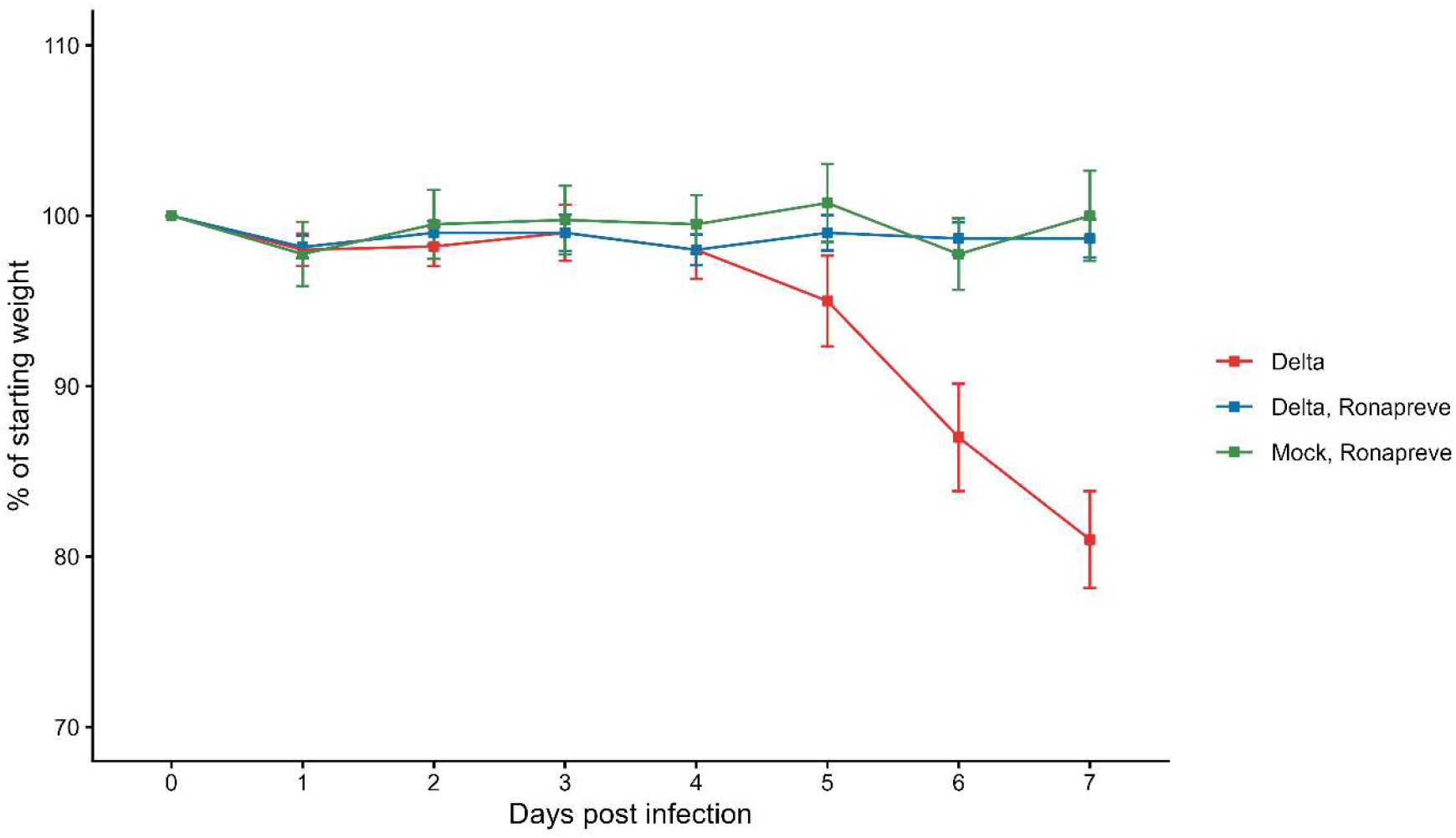
Body weights of K18-hACE2 mice after intranasal challenge with SARS-CoV-2 Delta at 10^3^ PFU, euthanized at 7 dpi. Each one cohort of 6 animals was treated with Ronapreve 24 hours post infection or remained untreated. Uninfected treated animals served as controls (data not shown). Weights are shown as % of body weight relative to the initial body weight at 0 dpi.

Ronapreve treatment via intraperitoneal injection of 400 µg Ronapreve at 24 hpi (Day 1) was associated with a slight drop in weight (approx. 2%) on the day of treatment, likely due to the peritoneal application procedure. The weight then increased slightly and remained almost stable in both infected and mock infected mice, albeit at an overall slightly lower level (98-99%) in the infected cohort. By 7 dpi, the body weight of the infected cohort was at 95-102% of the initial weight.

### The pharmacokinetic (PK) and biodistribution of Ronapreve in mock infected K18-hACE2 mice

Terminal serum and tissue concentrations of Ronapreve were evaluated in mock infected mice 6 days after treatment to assess whether SARS-CoV-2 infection alters the PK. The mice also served to determine if the therapeutic induces histological changes. Regarding the PK, differences were observed between the mAbs that constitute Ronapreve. Casirivimab concentrations were higher than imdevimab concentrations in both serum (16.5%, p = 0.03) and lung tissue (48.9%, p = 0.34), while concentrations in the spleen were comparable. In contrast, imdevimab concentrations were higher than casirivimab concentrations across all other tissues, by a mean of 38.4%. Specifically, imdevimab concentrations were significantly higher in the brain (70.2% higher; p = 0.03) compared to casirivimab (Figure 2a, b; Supplemental Table S6).

**Figure 2.**
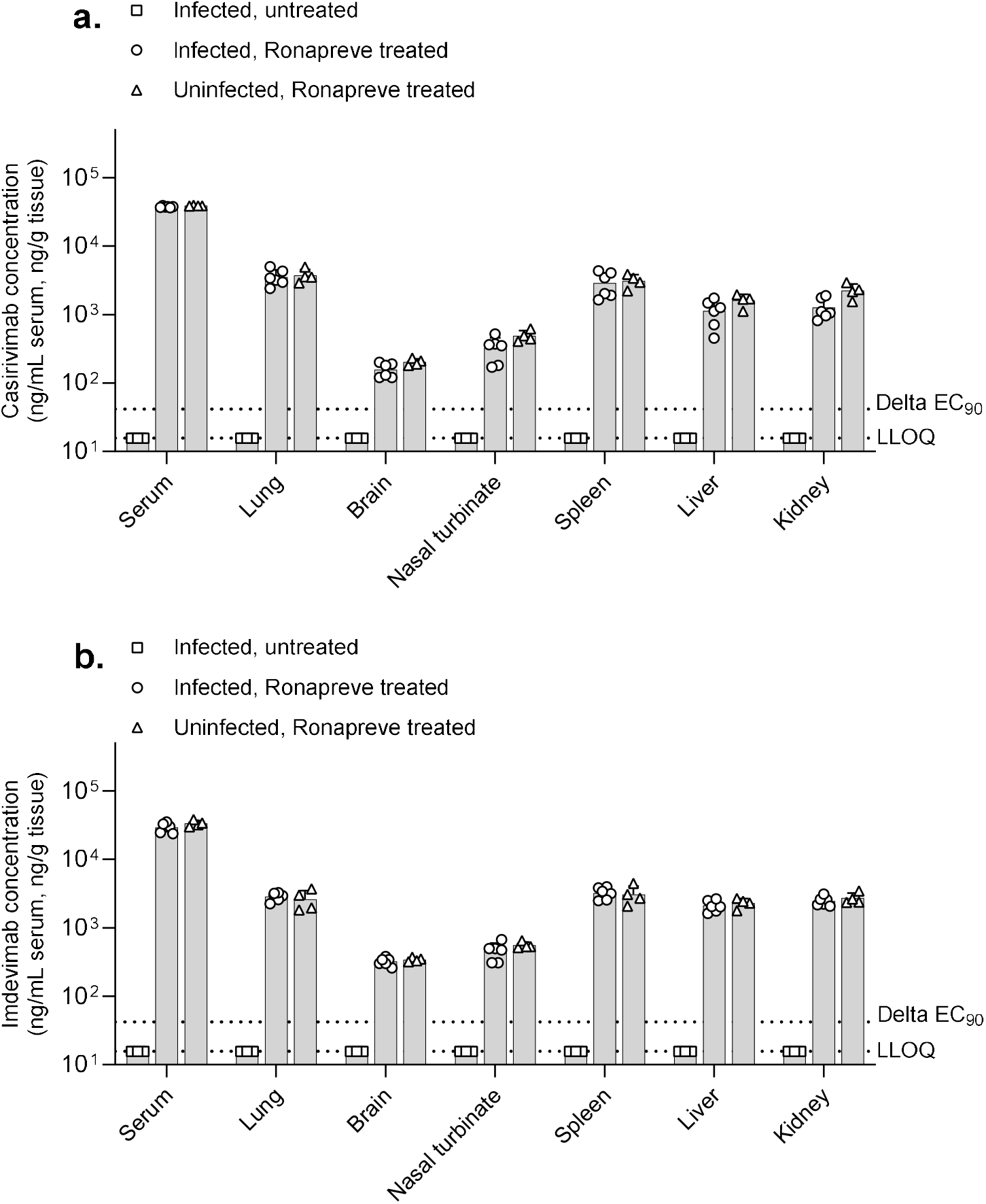
Concentrations of casirivimab (a.) and imdevimab (b.) in serum and tissues of K18-hACE2 mice at 7 dpi. Mice were intraperitoneally administered a single dose of Ronapreve (400 µg in saline) or saline alone (untreated cohort), 24 h after intranasal challenge with either SARS-CoV-2 Delta (10^3^ PFU in PBS) or PBS alone (uninfected cohort). Bars represent mean concentrations, with individual data points shown. SARS-CoV-2 Delta EC90 values are overlaid within the plots and were calculated from published EC50 values (*48*), assuming a Hill slope of 1. LLOQ: lower limit of quantification; EC90: concentration at which 90% inhibition of viral replication is observed.

Tissue-to-serum ratios were calculated to assess the extent of antibody distribution and to indicate tissue penetration relative to systemic exposure. Overall, the tissue penetration of both mAbs appeared low across the samples. Casirivimab showed the highest ratio in the lung (0.096), the ratio was lower for imdevimab (0.079). In the spleen, liver, and kidney, imdevimab concentrations exceeded those of casirivimab, with tissue-to-serum ratios of 0.095, 0.068, and 0.082, respectively, compared to 0.080, 0.041, and 0.057 for casirivimab. The penetration of mAbs was low in the brain and nasal turbinate compared to other tissues. Notably, imdevimab ratios in the brain were 2-fold higher than casirivimab (0.010 vs. 0.005), and higher ratios were also observed in the nasal turbinate (0.017 vs. 0.013). Overall, the results indicate distinct but overlapping tissue distribution patterns between the mAbs.

While ELISA-based quantification of tissue homogenates demonstrates mAb distribution in a tissue sample collected after the animal’s death, it cannot resolve whether the antibodies remain within the vasculature or have penetrated into the extravascular parenchyma. Therefore, to detect casirivimab and imdevimab *in situ*, an immunohistology protocol was established. This demonstrated the presence of both mAbs within vessel lumina and in the serum, with generally no evidence of their presence outside blood vessels (Figure 3).

**Figure 3.**
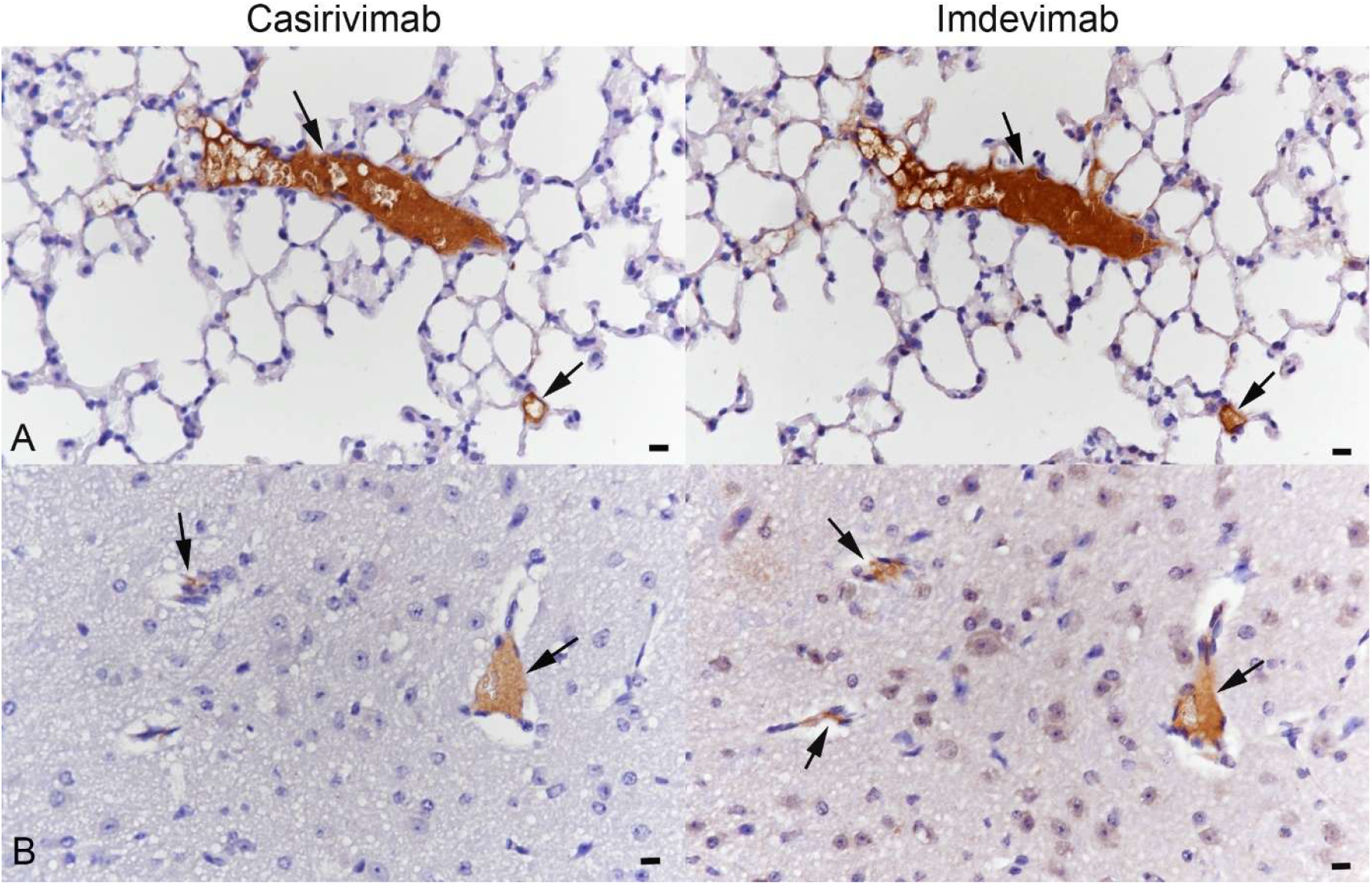
Ronapreve treated mock infected mouse. Both casirivimab and imdevimab are detected intravascularly, in the serum (arrows) but not outside vessels, in the parenchyma. **A)** Lung; **B)** Brain (cortex). Immunohistology, hematoxylin counterstain. Bars = 10 µm.

### Ronapreve treatment has minimal effects on the brain transcriptome and no effects on the brain metabolome and lipidome of mock infected mice

The transcriptomic analysis identified an outlier in the mock infected cohort (sample 3.3) which was excluded from all comparativeomic analyses. Based on the data of the remaining three animals (3.1, 3.2, 3.4), we found the Ronapreve treatment to be associated with minimal changes in the brain transcriptome, with significant up-regulation of only 6 genes: *Slc25a48, Snai3, Gm19963, Gpx6, Ido1*, and *Ggt6* (Figure 4A,B). GO enrichment of these genes (using ORA) show tabolome and lipidome of the mice (Figure 4C, D). ed pathways associated with immunity/inflammatory and metabolic biological processes, including T cell tolerance induction, chronic inflammatory response, kynurenine metabolic process, and quinolinate biosynthetic process (Figure 5). Their upregulation might be part of a direct response to the mAb boost.

**Figure 4.**
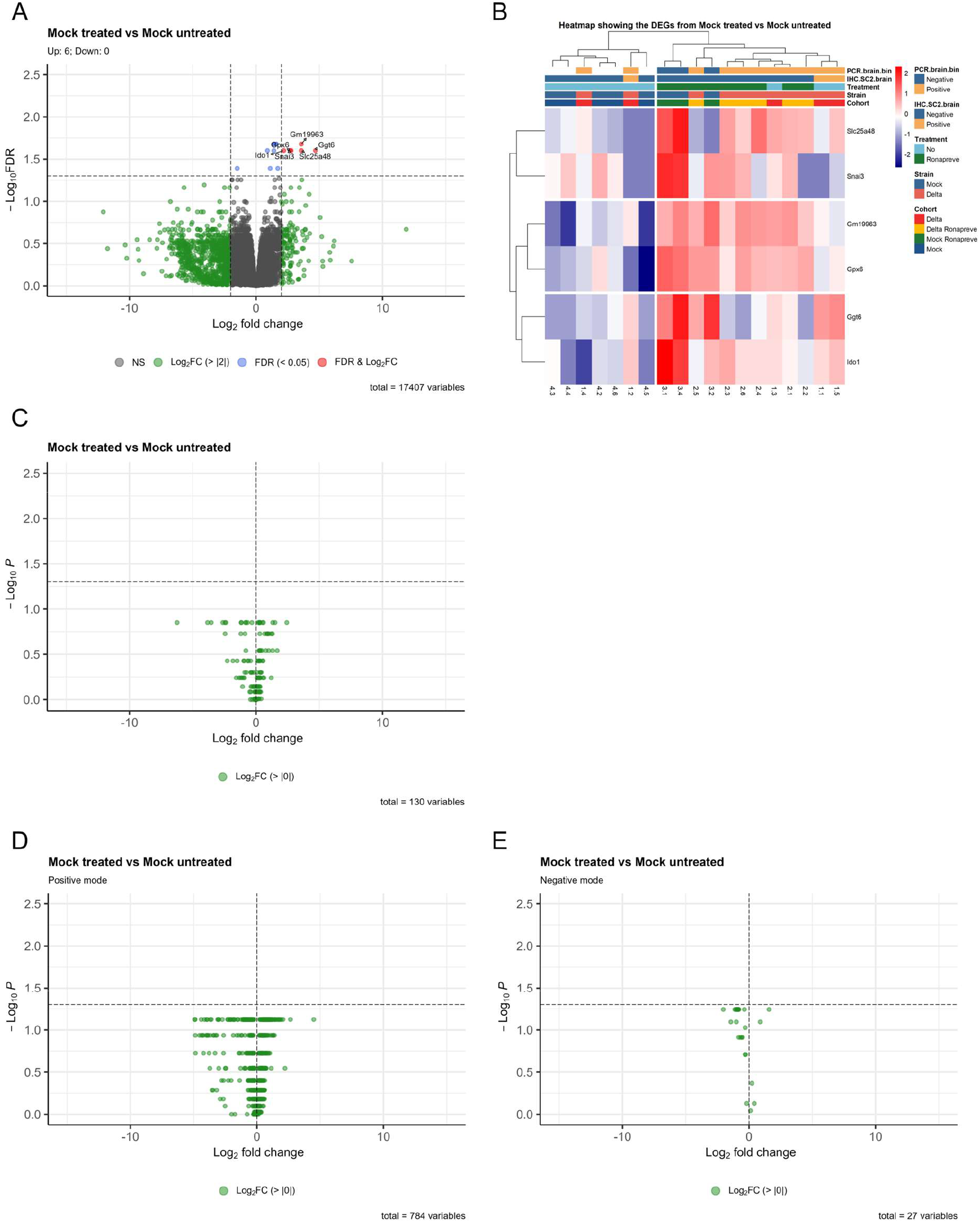
Multiomic analysis for the mock infected treated (mock treated) vs mock infected untreated (mock untreated) comparison pair. A,B) Transcriptomic analysis (thalamus). A) Volcano plot showing 6 significantly up-regulated genes in the mock infected treated cohort. B) Heatmap for the significantly up-regulated genes in the comparison pair. C) Metabolomic analysis (hypothalamus), volcano plot showing no significantly different metabolites. D) Lipidomic analysis (hypothalamus), volcano plots for the positive and negative modes, showing no significantly different lipids. DEGs: differentially expressed genes.

**Figure 5.**
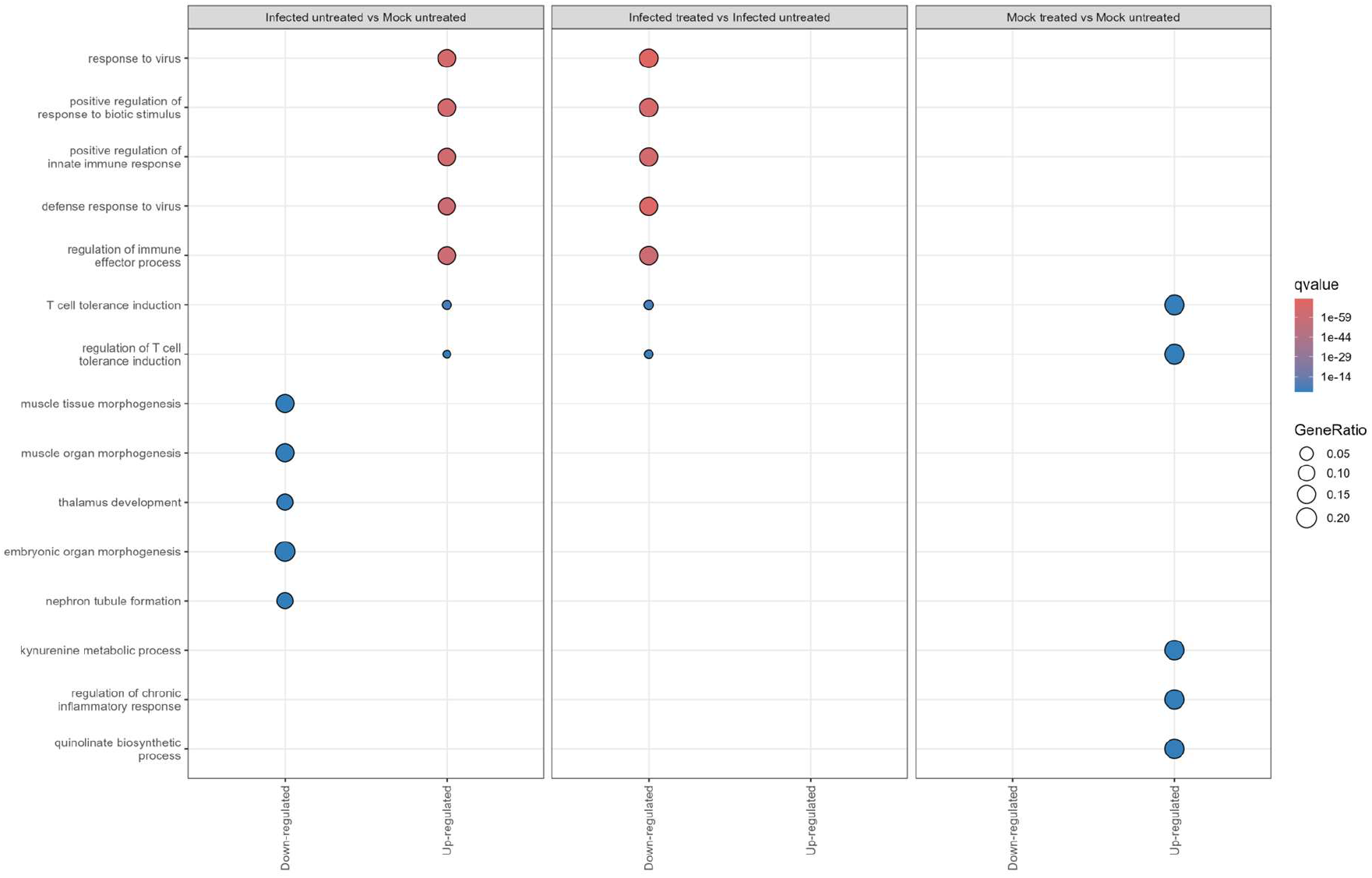
Thalamus, enrichment analysis with Gene Ontology (Biological Process) using the Over-Representation Analysis method. The top five (most significant) pathways for each comparison pair, and for each gene differential expression direction (i.e. up-regulated or down-regulated), are shown.

Ronapreve treatment was not associated with significant changes in the brain me

When the analyses were performed, including the outlier, this led to slightly more abundant changes in the brain transcriptome (significant up-regulation of 23 genes) (Supplemental Figure S2A,B). However, it introduced significant changes in the brain lipidome, with 130 lipids significantly more abundant and 106 lipids significantly less abundant in the treated mice (Supplemental Figure S2C,D). Enrichment analysis of the significant genes (ORA with GO) showed pathways associated with metabolic biological processes. The differentially expressed lipids covered various lipid classes, cellular components (endoplasmic reticulum, mitochondrion, plasma membrane, golgi apparatus, endosome/lysosome, lipid droplet) and functions (membrane component, lipid-mediated signalling, lipid storage) (Supplemental Figure S2E-H). Enrichment analysis on the lipids (Lipid Set Enrichment Analysis (LSEA) on the positive mode) confirmed enrichment of several cellular components, functions, lipid classifications, fatty acid properties, and physical or chemical properties (Supplemental Figure S2I). Amongst the top/bottom results were HexCer, sphingolipids, and glycosphingolipids (positively enriched) as well as triradylglycerols and triacylglycerols (negatively enriched). Additionally, positive enrichment of plasma membrane and endoplasmic reticulum, and negative enrichment of lipid droplet and lipid storage was found, the significance of which is unclear. Ultimately, however, the reason for such divergence between this sample and the rest of the cohort remains unknown.

### Post-challenge Ronapreve treatment of K18-hACE2 mice intranasally infected with SARS-CoV-2 Delta reduces virus burden and pathological effects in lungs and brain

We have previously shown in K18-hACE2 mouse cohorts examined at 6 and 9 days post SARS-CoV-2 Delta infection that the post-challenge Ronapreve treatment regimen applied in the present study reduces viral loads in the lungs; furthermore, we found convincing evidence that it also blocks viral brain infection (*20*). The present study investigated this finding further and comparatively examined cohorts of untreated and Ronapreve treated SARS-CoV-2 Delta infected mice (n=5 and 6, respectively) at 7 dpi.

Untreated mice exhibited high viral RNA loads in the **lungs** (>10^7^ copies of N1/µg of RNA relative to 18S; Supplemental Figure S3A). Immunohistology for viral NP showed that this was associated with viral antigen expression in patches of alveoli (type I and II pneumocytes). The histological examination confirmed the presence of the typical inflammatory processes observed in K18-hACE2 mice after SARS-CoV-2 VOC infection (*14*), represented by consolidated areas with activated type II pneumocytes, a few macrophages, several lymphocytes, rare neutrophils and scattered degenerate, occasionally desquamed cells (Supplemental Figure S4A).

After Ronapreve treatment, viral titres in the lungs did not exceed 6×10^5^ copies (range: 8.72×10^4^ – 6.33×10^5^ copies) and were thereby significantly reduced (Supplemental Figure S3A). Viral antigen expression was not detected by immunohistology, and the histological examination revealed limited changes, represented by small parenchymal leukocyte aggregates (lymphocytes, macrophages, rare neutrophils) and granulomatous infiltrates as well as a mild to moderate multifocal perivascular mononuclear, lymphocyte-dominated infiltration (Supplemental Figure S4B), similar to but less extensive than what we had observed in Ronapreve treated mice at 6 dpi (*20*). The detailed results in individual animals are provided in Supplemental Table S7.

All untreated mice harbored viral RNA in the **brain**, (Supplemental Table S7; Supplemental Figure S3B). Immunohistology showed widespread viral antigen expression in abundant neurons in the two brains with the highest viral RNA load (>2×10^10^ copies of N1/µg of RNA relative to 18S; animals 1.1 and 1.2; Supplemental Figure S5A), and was observed in clusters of positive neurons in the brain with 3.69×10^9^ copies of N1/µg of RNA relative to 18S (animal 1.5). In all three brains, this was accompanied by mild perivascular mononuclear infiltration (Supplemental Figure S5A) consistent with the mild non-suppurative encephalitis typically seen in K18-hACE2 mice infected with SARS-CoV-2 VOC at this time point (*13, 14*). The remaining two brains, which harboured <6×10^7^ copies of N1/µg of RNA relative to 18S, viral antigen was not detected, and there was no evidence of any histological changes. The detailed results in individual animals are provided in Supplemental Table S7.

The Ronapreve treated mice harboured viral RNA at very low levels in the brain (2.25×10^4^ to 1.45×10^5^ copies of N1/µg of RNA relative to 18S; Supplemental Figure S3B), without evidence of viral antigen or viral RNA signal, as shown by immunohistology and RNA-ISH, and without any histological changes (Supplemental Figure S5B), in line with the results of our previous study and confirming that systemic Ronpreve treatment blocked viral spread to the brain (*20*). The detailed results in individual animals are provided in Supplemental Table S7.

### The PK and biodistribution of Ronapreve in SARS-CoV-2 Delta-infected K18-hACE2 mice

Tissue and serum concentrations of casirivimab and imdevimab showed distribution profiles similar to those observed in uninfected animals. Serum concentrations were slightly lower following infection (casirivimab: −4.5%; imdevimab: −10.3%), but this did not correspond to marked changes in tissue distribution. Tissue-to-serum ratios remained low across all organs, and the same relative trends were retained, with the lungs exhibiting the highest ratios and the brains the lowest (Figure 2; Supplemental Table S6). Imdevimab was consistently detected at higher concentrations in tissues than casirivimab in most organs, with notable differences in the brain, suggesting higher tissue penetration (Figure 2). The results suggest that SARS-CoV-2 infection does not substantially alter the biodistribution of Ronapreve. The calculated EC_90_ values (*48*) for both mAbs are shown in Figure 2a and b and illustrate that all mouse serum and tissue concentrations at 7 dpi exceeded the EC_90_ threshold for the Delta variant by a substantial margin. In infected animals, casirivimab concentrations were approximately 900-fold higher in serum, more than 88-fold higher in lung, and over 3.8-fold higher in brain compared to the calculated EC_90_. Similarly, imdevimab concentrations were approximately 700-fold higher in serum, more than 66-fold higher in lung, and over 7.5-fold higher in brain relative to the calculated EC_90_.

Immunohistology for casirivimab and imdevimab in the tissues demonstrated the presence of both monoclonal antibodies within vessel lumina, in the serum (Figure 6). In the lung, in association with areas harbouring an inflammatory infiltration, there was occasional weak staining of macrophages in affected areas (Figure 6B). In the brain of treated animals, where neither viral antigen nor an inflammatory response was observed, there was no evidence of casirivimab or imdevimab expression outside blood vessels (Figure 6D).

**Figure 6.**
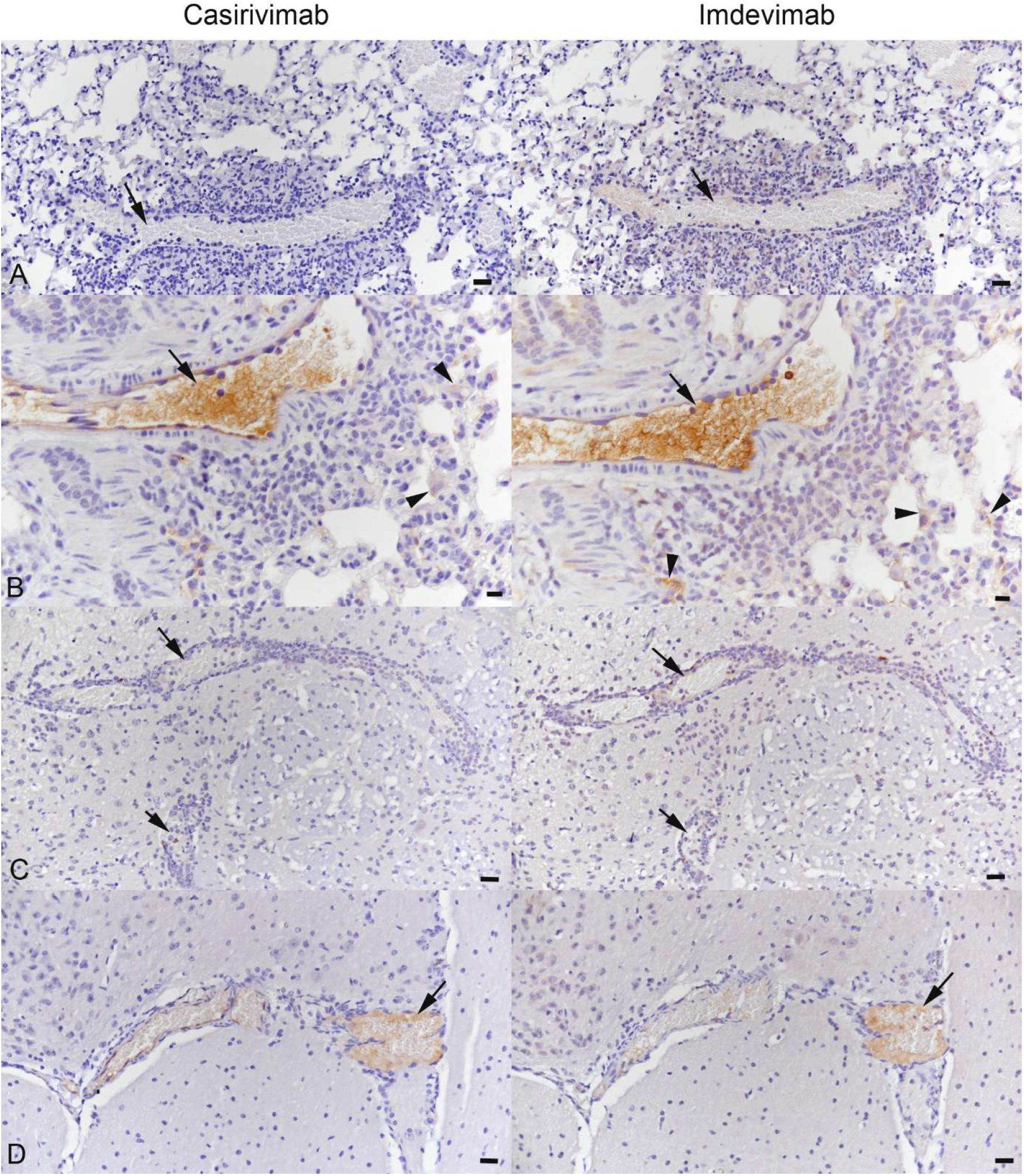
Lung (A, B) and brain (C, D) of K18-hACE2 mice examined 7 days after intranasal challenge with SARS-CoV-2 Delta at 10^3^ PFU, with and without treatment with Ronapreve. **A, B)** Lung. Vessels (arrows) with perivascular leukocyte infiltrate. In the untreated animal (A), staining for casirivimab and imdevimab shows no reaction, neither in the serum nor in the surrounding parenchyma. In the treated animal (B), a strong serum reaction is seen for both monoclonal antibodies. There are also a few macrophages in adjacent alveoli that exhibit a weak focal cytoplasmic reaction (arrowheads), indicating serum uptake after serum leakage from the vessel. **C, D)** Brain. In the untreated animal (**C**), viral infection of the brainstem (confirmed by PCR and immunohistology for viral NP) is associated with moderate leukocyte recruitment and perivascular accumulation, as previously reported (*13, 14*). In the treated animal (**D**; negative for SARS-CoV-2 by PCR and immunohistology), the serum stains positive for both monoclonal antibodies. There is no evidence of an inflammatory reaction. The arrows highlight veins in the brain parenchyma. Immunohistology, hematoxylin counterstain.

### The minimal viral RNA burden in the brain of Ronapreve treated infected mice is associated with very limited alterations of the brain transcriptome and no significant changes in metabolome and lipidome

As previously shown by our group (*14*), brain infection with neuronal viral antigen expression and morphologically apparent inflammatory infiltration after intranasal challenge with SARS-CoV-2 Delta was associated with distinct transcriptomic profiles, characterised by an inflammatory and antiviral/immune response (Figure 5; Figure 7A, B). However, in the present study which also included brains of untreated mice without overt viral antigen expression, as shown by immunohistology, and/or histological changes, we detected evidence of correlation of the transcriptomic profiles with both the presence of viral NP in the tissue and the viral RNA load in the brain, as measured by qRT-PCR (Figure 7C, D). Indeed, we found the infected untreated brains to form a continuum, with higher viral loads associated with more pronounced transcriptional changes; brains positive for viral NP in neurons were clearly separated from the NP-negative ones and, interestingly, the infected brain with the lowest viral RNA load (and no evidence of viral antigen expression) was overlapping with the mock-infected cohort (Figure 7C). Unsurprisingly, infected animals with higher viral loads in the lung and brain and SARS-CoV-2 NP expression showed a strong correlation between these parameters and PC1 (Figure 7D).

**Figure 7.**
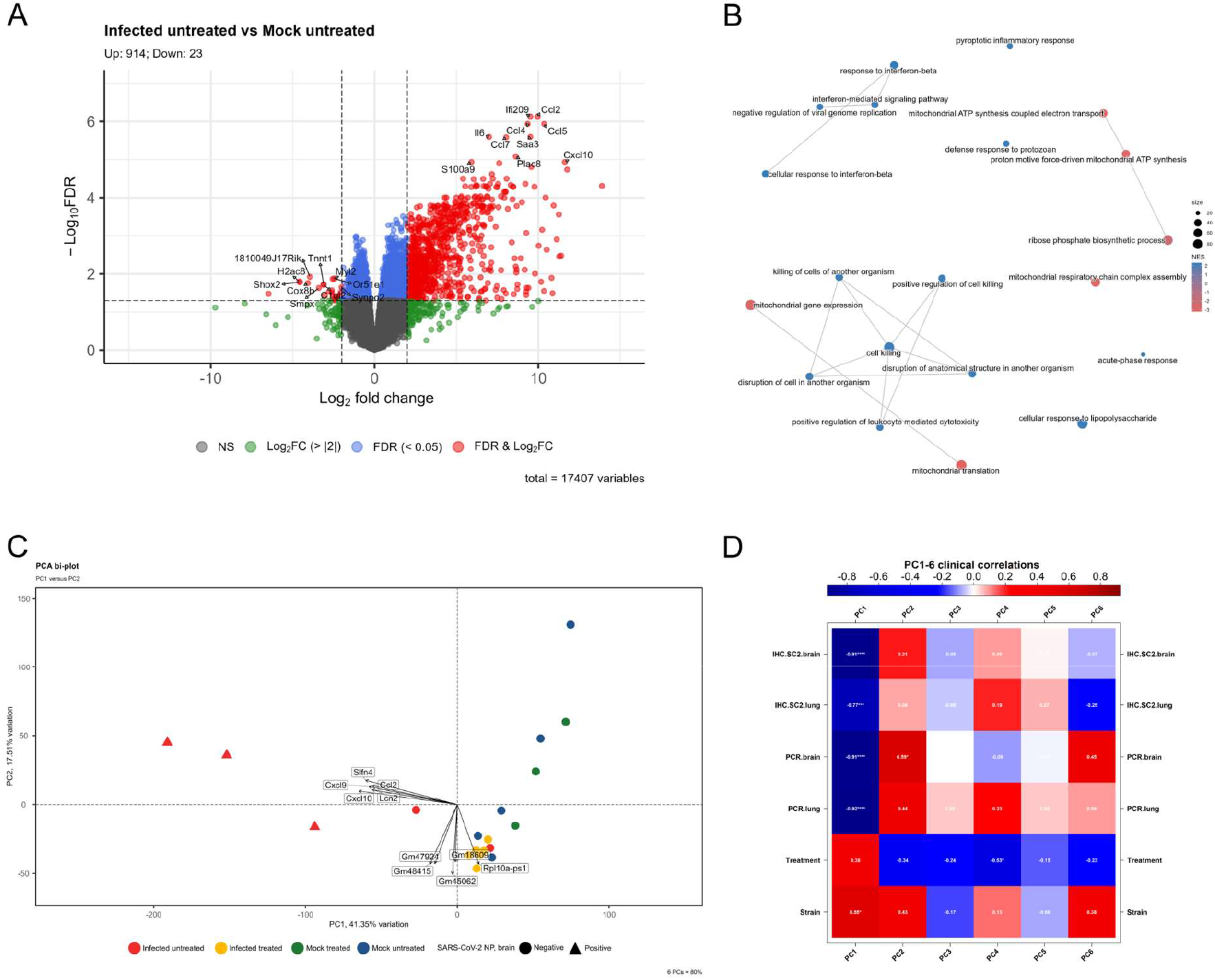
Transcriptomic analysis (thalamus) for the infected untreated vs mock infected untreated (mock untreated) comparison pair. A) Volcano plot showing that most differentially expressed genes are up-regulated in the infected untreated cohort. B) Gene Set Enrichment Analysis (GSEA) using Gene Ontology (GO) and visualised after applying clusterProfiler simplify() function to remove redundant GO terms; the top 20 terms are shown. C) Principal Component Analysis (PCA) showing the clustering of the four cohorts. D) Eigencorplot showing the correlation between several clinical/morphological parameters and the Principal Components (PCs). FDR: False Discovery Rate; IHC.SC2.brain: IH against SARS-CoV-2 NP in the brain; IHC.SC2.lung: IH against SARS-CoV-2 NP in the lung; PCR.brain: qRT-PCR for SARS-CoV-2 in the brain; PCR.lung: qRT-PCR for SARS-CoV-2 in the lung; Strain: virus-related experimental design (i.e. SARS-CoV-2 Delta infected or mock infected); Treatment: cohort with or without Ronapreve treatment.

Investigation into the brain metabolome showed that SARS-CoV-2 infection of the brain was associated with significantly higher abundance of 9 metabolites and lower abundance of 16 metabolites (Figure 8C, D). Although the infected untreated brains (independent of the detection of viral NP) tended to cluster separately, this was less clear for the metabolome than for the transcriptome (Figure 8A). Indeed, correlation between lung and brain viral loads and SARS-CoV-2 NP expression was observed only with PC2 and PC4, which allowed better separation of the infected untreated cohort from the other cohorts (Figure 8B,E).

**Figure 8.**
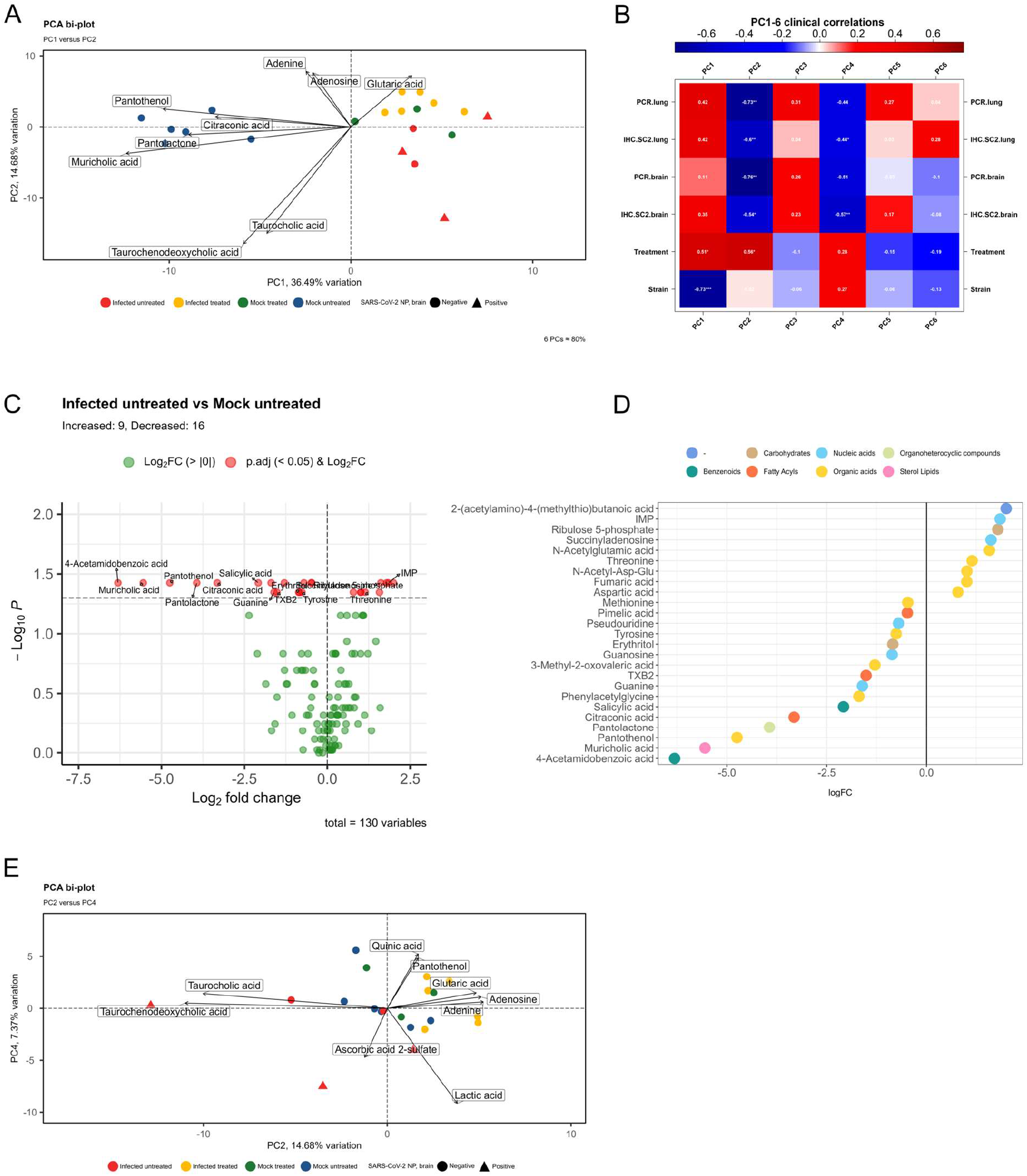
Metabolomic analysis (hypothalamus). A) Principal Component Analysis (PCA) showing the clustering of the four cohorts. B) Eigencorplot showing the correlation between several clinical/morphological parameters and the PCs. C) Volcano plot showing the significantly differential metabolites for the infected untreated vs mock infected untreated (mock untreated) comparison pair. D) The differential metabolites are colour-coded based on their super class. E) Principal Component Analysis with Principal Components 2 and 4 (PC2, PC4) to achieve better separation of the infected untreated cohort. IHC.SC2.brain: IH against SARS-CoV-2 NP in the brain; IHC.SC2.lung: IH against SARS-CoV-2 NP in the lung; Log_2_FC/LogFC: Log_2_ Fold Change; P: adjusted P-value; PCR.brain: qRT-PCR for SARS-CoV-2 in the brain; PCR.lung: qRT-PCR for SARS-CoV-2 in the lung; Strain: virus-related experimental design (i.e. SARS-CoV-2 Delta infected or mock-infected); Treatment: cohort with or without Ronapreve treatment.

Additionally, we showed that brain infection was not associated with significant changes in the brain lipidome (Figure 9A, B). Some correlation between lung and brain viral loads and SARS-CoV-2 NP expression was observed with PC2, PC5, PC6, and PC7 and indeed, better separation of all infected brains was achieved when considering these principal components (Figure 9C-I).

**Figure 9.**
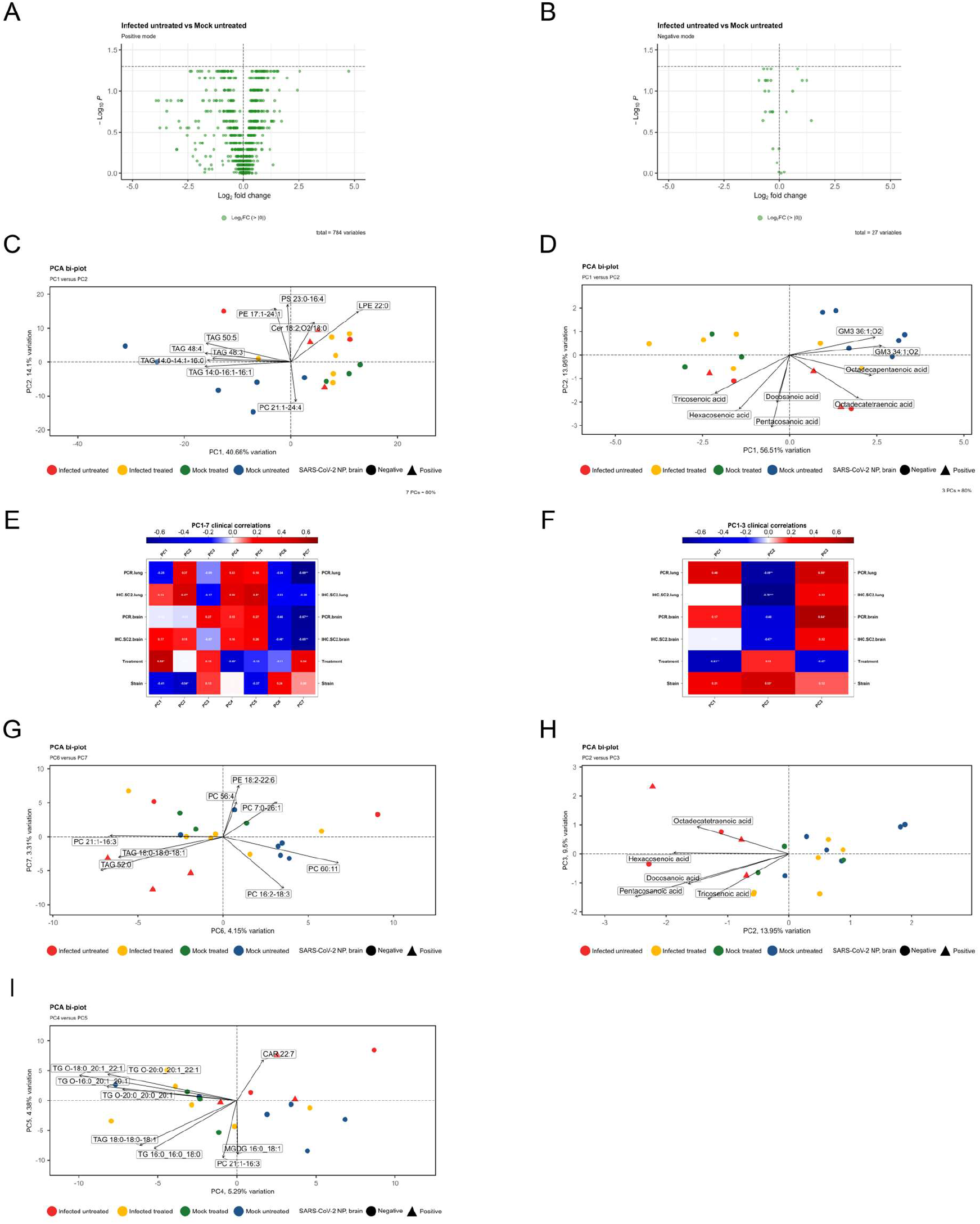
Lipidomic analysis (hypothalamus). A,B) Volcano plots for the positive (A) and negative (B) modes, for the infected untreated – mock infected untreated (mock untreated) comparison pair, showing no significant lipids. C,D) Principal Component Analysis (PCA) for the positive (C) and negative (D) modes showing the clustering of the cohorts. E,F) Eigencorplot, positive (E) and negative (F) modes, showing the correlation between several clinical/morphological parameters and the PCs. G-I) Principal Component Analysis with different combinations of Principal Components (PCs) to achieve better separation of the infected untreated cohort, both for the positive (G, I) and negative (H) modes. IHC.SC2.brain: IH against SARS-CoV-2 NP in the brain; IHC.SC2.lung: IH against SARS-CoV-2 NP in the lung; Log_2_FC: Log_2_ Fold Change; P: adjusted P-value; PCR.brain: qRT-PCR for SARS-CoV-2 in the brain; PCR.lung: qRT-PCR for SARS-CoV-2 in the lung; Strain: virus-related experimental design (i.e. SARS-CoV-2 Delta infected or mock-infected); Treatment: cohort with or without Ronapreve treatment.

The blocking of viral spread to the brain, i.e. the effect of Ronapreve treatment after intranasal challenge of the mice with SARS-CoV-2 Delta, was also observed at the transcriptome level. The majority of the infected untreated brains and the two mock infected (with and without Ronapreve treatment) cohorts clustered separately, forming the two extremes of a spectrum (infected vs non-infected), although with noticeable interindividual variation within each cohort. The brains of the infected Ronapreve treated mice fell between those two extremes, despite some overlap with both the mock infected cohorts and the infected brain with the lowest viral load from the untreated group (Figure 7C). This is confirmed by the differential gene expression which showed the limited brain response in the Ronapreve treated infected cohort. Comparison of the infected Ronapreve treated vs untreated pair confirmed the strong up-regulation of genes in the latter cohort (Figure 10A), with enrichment of pathways associated with antiviral/immune response (Figure 5), similar to the comparison of the infected vs mock infected untreated groups (Figure 5, Figure 7A). The infected, treated and untreated cohorts showed no significantly differences when the effect of Ronapreve treatment was removed, revealing the consequence of the very limited infectious burden in the brain on gene expression following treatment (Figure 10B). The direct comparison between the brains of infected and mock infected Ronapreve treated mice showed no significant differences between these two cohorts, confirming the effect of reduced viral burden secondary to Ronapreve treatment on the brain transcriptome (Figure 10C).

**Figure 10.**
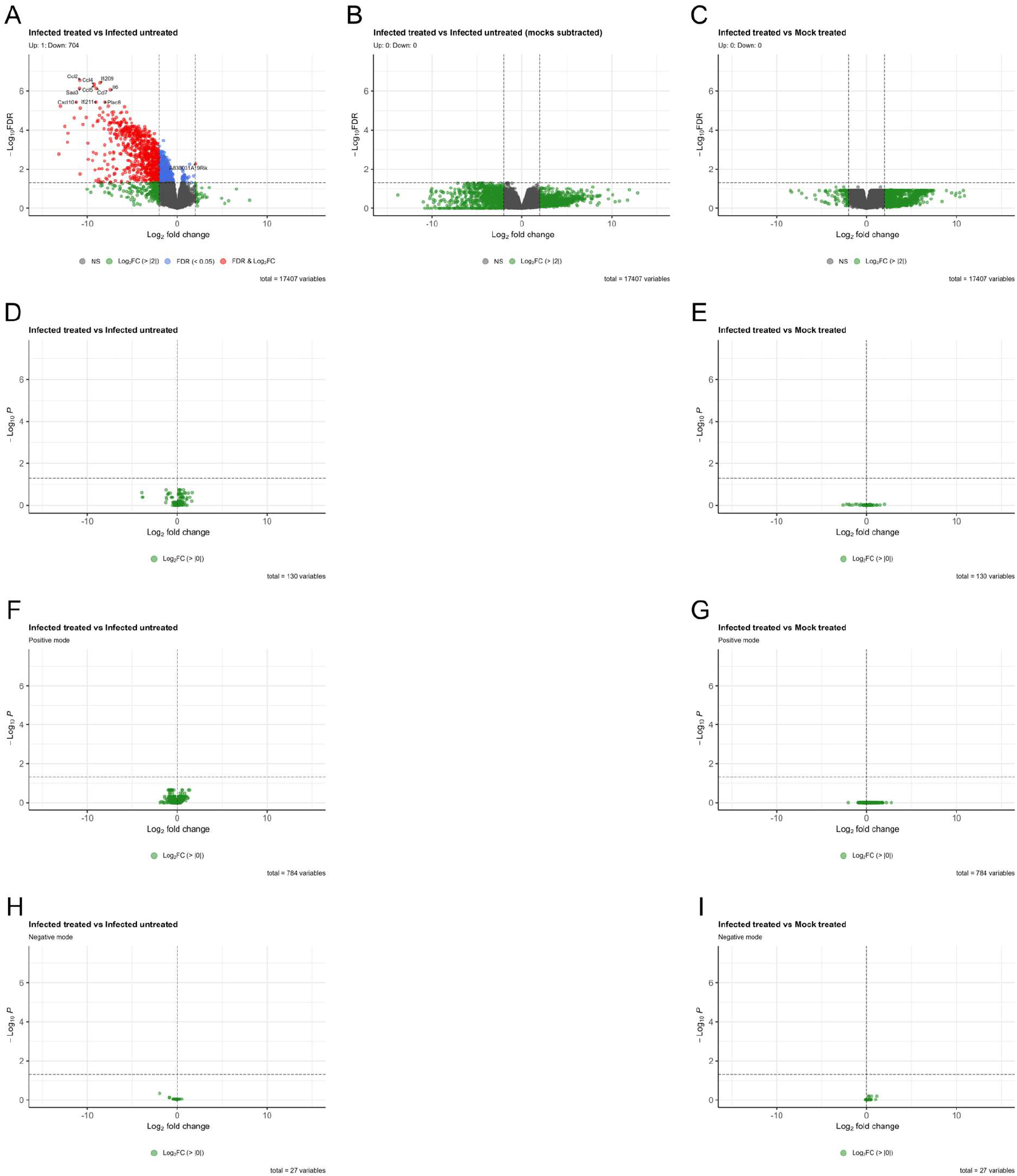
Volcano plots for the different comparaison pairs for A-C) Transcriptomic (thalamus). D-E) Metabolomic (hypothalamus). F-G) Lipidomic, positive mode (hypothalamus). H-I) Lipidomic, negative mode (hypothalamus). FDR: False Discovery Rate; Log_2_FC: Log_2_ Fold Change; Mock treated: mock infected treated; Mock untreated: mock infected untreated; P: adjusted P-value.

Interestingly, the brain metabolome and lipidome of the infected Ronapreve treated cohort showed no significant difference to its untreated counterpart, or to the mock infected treated cohort (Figure 10D-I).

## Discussion

This study provides mechanistic insight into the processes that prevent SARS-CoV-2 neuroinvasion in K18-hACE2 mice after systemic post-challenge administration of Ronapreve. PK analysis demonstrated that both casirivimab and imdevimab achieved high systemic concentrations at 7 dpi, with no significant difference between infected and mock infected animals. Brain and nasal turbinate mAb concentrations were significantly lower, with tissue-to-serum ratios that were the lowest among all tissues analysed. However, concentrations of both mAbs were substantially above the calculated EC_90_ for SARS-CoV-2 Delta (*48*) in homogenised samples at 7 dpi. Immunohistology confirmed that both mAbs remained confined to the brain vasculature, with no evidence of extravasation into the parenchyma. These findings are consistent with the well-established restriction of IgG transport across an intact BBB, attributed to the high molecular weight (∼150 kDa) and hydrophilic nature of IgG molecules (*28, 49*). Interestingly, while the two mAbs clearly did not reach the brain parenchyma, minimal changes in the brain transcriptome were generally observed in healthy treated mice 6 days after mAb application, though they did not suggest a specific tissue response. The fact that one animal in this cohort presented as an outlier, with pronounced, yet equally nonspecific, lipidomic changes, could reflect interindividual variability in the response to the mAbs.

Despite the lack of mAb extravasation, only very low quantities of viral RNA and neither viral antigen nor histological changes were detected in the brains of the intranasally infected Ronapreve treated mice, providing strong evidence of a mechanism in which neuroinvasion is blocked at peripheral sites rather than within the CNS itself. The effect of the treatment on brain infection was further confirmed at the transcriptome level, as the brain transcriptome of the infected Ronapreve treated cohort showed no significantly different genes compared to the mock infected treated cohort. Additionally, no changes were observed between these two cohorts at the metabolomic and lipidomic level.

In the lungs, while both mAbs were predominantly confined to the vasculature, limited extravasation into alveolar spaces was evident in areas of inflammation, where individual macrophages exhibited immunoreactivity for both antibodies. This is consistent with our previous findings suggesting that antibody-opsonised virus was phagocytosed by macrophages and hence confined within granulomatous infiltrates (*20*). Viral antigen was not detected in the lung at 7 dpi, and low quantities of viral RNA were identified in treated animals. Therapeutic mAbs typically extravasate into the lung parenchyma, facilitated by neonatal Fc receptor (FcRn) transport and vascular permeability, to neutralise the virus within the alveolar space (*50, 51*). The detection of antibodies within alveolar macrophages and high mAb concentrations in homogenised lung samples in the present study indicates that sufficient antibody reached the alveolar compartment to facilitate viral clearance. Of note, terminal blood collection prior to tissue harvest may have reduced residual intravascular antibody levels in tissue homogenates, potentially underestimating the contribution of intravascular mAbs to measured tissue concentrations.

Both mAbs were detected in homogenised nasal turbinate samples. While these measurements include intravascular antibody and cannot directly assess mucosal surface concentrations, previous studies have shown that systemically administered IgG enters the nasal lining fluid following infection (*52*) and vaccination (*53*). This mechanism likely mediates the presence of neutralising mAbs at the olfactory epithelium. Similarly, pharmacologically active concentrations of tixagevimab and cilgavimab (Evusheld) have been demonstrated in nasal lining fluid following intramuscular administration, reaching ∼1-2% of serum concentrations, roughly 40-fold above the IC_50_, and persisting for at least 30 days (*54, 55*). The presence of neutralising mAbs at the olfactory epithelium likely reduced infection of and replication in the epithelial cells and prevented infection of olfactory sensory neurons, thereby impairing the viral entry required for subsequent retrograde transport, which underlies CNS dissemination in our model (*15, 20*). Given the prolonged half-life of Ronapreve (∼25-37 days in humans (*2, 56*)), a sustained exposure at mucosal surfaces likely contributes to prolonged antiviral activity. Importantly, mAb treatment at 24 hpi markedly reduced CNS infection, preventing neuronal viral antigen accumulation and the associated inflammatory response, despite active viral replication in respiratory tissues. This demonstrates a viable therapeutic window for preventing CNS infection. The protective effect aligns with the neuroinvasion timeline in this model (typically 3-5 dpi (*15, 20*)), indicating that early intervention can block spread to the brain. Future studies should determine how late treatment can be delayed while maintaining neuroprotection. Of note, this study examined antibody distribution at a single terminal time point, thereby limiting the assessment of time-dependent changes in tissue distribution or the determination of peak tissue concentrations.

Modest differences in the distribution of casirivimab and imdevimab were observed: imdevimab achieved higher tissue-to-serum ratios across several tissues, including the brain and nasal turbinates, whereas casirivimab ratios were higher in the lung. Such differences likely reflect intrinsic mAb molecular properties influencing FcRn binding, isoelectric point, or nonspecific tissue interactions (*57, 58*).

The findings of the present study have relevance to other respiratory viruses with the potential for neuroinvasion. Several influenza A virus strains exhibit olfactory neurotropism, accessing the CNS by infecting the olfactory epithelium and subsequently spreading along the olfactory nerve into the olfactory bulb (*24*). Similarly, RSV, although only rarely neuroinvasive in humans, has been shown in murine models to access the CNS following intranasal infection via the olfactory route and subsequent spread to the brain (*59*). The olfactory route is also used by neuropathic viruses, such as rabies and Japanese encephalitis virus, as a route of entry to the brain (*24*). Together, these parallels underscore that this route represents a shared anatomical vulnerability across multiple respiratory pathogens and highlight the therapeutic relevance of antiviral agents that restrict viral replication at this site, even in the absence of BBB penetration.

These data suggest that systemic administration of neutralising antibodies prevents SARS-CoV-2 neuroinvasion by efficiently restricting viral replication at the nasal mucosa, without requiring BBB penetration. Persistent neurological symptoms, including cognitive impairment (‘brain fog’) and chronic fatigue, are hallmarks of Long-COVID (*60*). While direct viral neuroinvasion has been demonstrated in severe human cases and animal models (*13, 20, 21*), the extent to which it contributes to Long-COVID neurological symptoms remains debated (*61, 62*). Nevertheless, early intervention with neutralising antibodies that block olfactory-mediated CNS entry may reduce the risk of potential neurological sequelae. Notably, while some studies in human COVID-19 patients have detected SARS-CoV-2 in olfactory structures (*21*), the predominant pattern appears to involve infection of sustentacular cells with minimal infection of olfactory neurons and limited neuroinvasion into the olfactory bulb (*62–64*), suggesting species-specific differences in neuroinvasive potential. Given that multiple respiratory viruses utilise the olfactory route for entry into the brain, these findings could have broader implications for the development of antiviral biologics and small drug molecules against future emerging pathogens with neuroinvasive potential.

## Ethics approval

Animal work was approved by the local University of Liverpool Animal Welfare and Ethical Review Body and performed under UK Home Office Project Licence PP4715265. It was undertaken in accordance with locally approved risk assessments and standard operating procedures.

## Authors’ contribution

AK (Anja Kipar), JPS, AO: conception and design of the project. JAS, SDN, RPR, AK (Adam Kirby), PS, LT, PC: methodology. PS, AK (Adam Kirby): in vivo experimental work. LT, JH, EGT, EK, JS: pharmacokinetic analyses. JAS, SDN, RPR, PS, AK (Adam Kirby), LT, PC: data collection. JAS, SDN, RPR, LT, AK (Anja Kipar): data analysis. JAS, SDN, AK (Anja Kipar), LT: writing of the original draft. JAS, SDN, AK (Anja Kipar), RPR, LT: review and editing. AK (Anja Kipar), JPS, AO: funding acquisition. AK (Anja Kipar): supervision of the project. All authors read and approved the manuscript.

## Acknowledgements

We are grateful to the technical staff in the Histology Laboratory, Institute of Veterinary Pathology, Vetsuisse Faculty, University of Zurich (IVPZ), for excellent technical support. Also, we thank the technical and scientific staff of the Functional Genomics Center Zurich (FGCZ), including Dr Alaa Othman, Dr Martina Zanella, and Dr Michelle Reid, ETH Zurich, University of Zurich, for their support regarding the acquisition and the interpretation of the metabolomic and lipidomic datasets. We would like to thank Professor Wei-Chung Cheng and Dr. Pei-Chun Shen at Program for Cancer Molecular Biology and Drug Discovery, College of Medicine, China Medical University, for their help in implementing their R package, LipidSigR, to our analysis.

