## Supplemental Material for "The biodistribution and effect of post-exposure neutralising monoclonal antibody treatment in a mouse model of SARS-CoV-2 infection with viral spread to the brain"

**Supplemental Table S1.** Summary of the experimental design and of the mice cohorts used in the study.

| **Cohort designation** | **No of mice** | **SARS-CoV-2 Delta infection** | **Ronapreve treatment** |
| --- | --- | --- | --- |
| Infected untreated | 5 | Y | N |
| Infected treated | 6 | Y | Y |
| Mock infected treated | 4^a^ | N | Y |
| Mock infected untreated | 6^b^ | N | N |

^a^ The transcriptomic analysis showed animal 3.3 as a clear outlier in this cohort. All -omics analyses were run without this sample (the results of the transcriptomic and lipidomic analyses with this animal are shown in Supplemental Figure S2).

^b^ For the transcriptomic analysis, animal 4.1 was excluded as it showed as an outlier on the Principal Component Analysis.

**Supplemental Table S2.** Details of the comparisons and contrasts used for the single -omics analyses.

A) Transcriptomic analysis

| **Comparison** | **limma::makeContrasts()** | **Transcriptional changes investigated** |
| --- | --- | --- |
| Infected untreated vs Mock infected untreated | Infected_untreated–Mock_untreated | Effect of SARS-CoV-2 Delta infection in untreated mice |
| Mock infected treated vs Mock infected untreated | Mock_treated-Mock_untreated | Effect of Ronapreve treatment in mock infected mice |
| Infected treated vs Infected untreated | Infected_treated-Infected_untreated | Effect of Ronapreve treatment in SARS-CoV-2 Delta infected mice |
| Infected treated vs Infected untreated (subtraction of the respective mock cohorts) | (Infected_treated-Mock_treated)-( Infected_untreated–Mock_untreated) | Effect of Ronapreve treatment in SARS-CoV-2 Delta infected mice |
| Infected treated vs Mock infected treated | Infected_treated-Mock_treated | Effect of SARS-CoV-2 Delta infection in Ronapreve treated mice |

B) Metabolomic and lipidomic analyses

| **Comparison** | **Metabolic changes investigated** |
| --- | --- |
| Infected untreated vs Mock infected untreated | Effect of SARS-CoV-2 Delta infection |
| Mock infected treated vs Mock infected untreated | Effect of Ronapreve treatment in mock infected mice |
| Infected treated vs Infected untreated | Effect of the Ronapreve treatment in SARS-CoV-2 Delta infected mice |
| Infected treated vs Mock infected treated | Effect of SARS-CoV-2 Delta infection in Ronapreve treated mice |

**Supplemental Table S3.** List of the R packages used for the -omics analyses.

A) Transcriptomic analysis

| **Package** | **Version** | **Reference** |
| --- | --- | --- |
| tximport | 1.32.0 | (*1*) |
| edgeR | 4.8.2 | (*2*) |
| PCAtools | 2.22.1 | (*3*) |
| limma | 3.66.0 | (*4*) |
| EnhancedVolcano | 1.28.2 | (*5*) |
| clusterProfiler | 4.18.4 | (*6–9*) |
| fgsea | 1.36.2 | (*10*) |
| AnnotationDbi | 1.72.0 | (*11*) |
| org.Mm.eg.db | 3.22.0 | (*12*) |
| ggplot2 | 4.0.1 | (*13*) |
| dplyr | 1.1.4 | (*14*) |
| patchwork | 1.3.2 | (*15*) |
| stringr | 1.6.0 | (*16*) |
| UpSetR | 1.4.0 | (*17*) |
| tibble | 3.3.1 | (*18*) |
| pheatmap | 1.0.13 | (*19*) |
| limma | 3.66.0 | (*4*) |
| enrichplot | 1.30.4 | (*20*) |

B) Metabolomic analysis

| **Package** | **Version** | **Reference** |
| --- | --- | --- |
| tidymass | 2.0.10 | (*21*) |
| massdataset | 0.99.0 | (*21*) |
| masscleaner | 1.0.12 | (*21*) |
| massqc | 1.0.8 | (*21*) |
| massstat | 1.0.6 | (*21*) |
| ggplot2 | 4.0.1 | (*13*) |
| rstatix | 0.7.3 | (*22*) |
| EnhancedVolcano | 1.28.2 | (*5*) |
| dplyr | 1.1.4 | (*14*) |
| tidyr | 1.3.2 | (*23*) |
| stringr | 1.6.0 | (*16*) |
| patchwork | 1.3.2 | (*15*) |
| PCAtools | 2.22.1 | (*3*) |
| RefMet | 1.0.0 | (*24*) |

C) Lipidomic analysis

| **Package** | **Version** | **References** |
| --- | --- | --- |
| LipidSigR | 1.0.4 | (*25–27*) |
| rgoslin | 1.14.0 | (*28, 29*) |
| RefMet | 1.0.0 | (*24*) |
| SummarizedExperiment | 1.40.0 | (*30*) |
| PCAtools | 2.22.1 | (*3*) |
| dplyr | 1.1.4 | (*14*) |
| stringr | 1.6.0 | (*16*) |
| tidyr | 1.3.2 | (*23*) |
| EnhancedVolcano | 1.28.2 | (*5*) |
| ggplot2 | 4.0.1 | (*13*) |
| patchwork | 1.3.2 | (*15*) |
| rstatix | 0.7.3 | (*22*) |
| tibble | 3.3.1 | (*18*) |

**Supplemental Table S4.** List of the R packages used for weight curves and the qRT-PCR results.

| **Package** | **Version** | **Reference** |
| --- | --- | --- |
| Rmisc | 1.5.1 | (*31*) |
| ggplot2 | 4.0.0 | (*13*) |
| rstatix | 0.7.2 | (*22*) |
| ggpubr | 0.6.1 | (*32*) |
| tidyr | 1.3.1 | (*23*) |
| dplyr | 1.1.4 | (*14*) |
| patchwork | 1.3.2 | (*15*) |

**Supplemental Table S5.** Statistical results for the qRT-PCR results of the lung and the brain, comparing SARS-CoV-2 Delta-infected untreated mice and SARS-CoV-2 Delta-infected, Ronapreve-treated mice.

Shapiro-Wilk test

| **Group** | **Organ** | **Statistic** | **p-value** |
| --- | --- | --- | --- |
| Infected untreated | PCR.brain | 0.7805871 | 0.05576359 |
| Infected untreated | PCR.lung | 0.9669397 | 0.85527247 |
| Infected treated | PCR.brain | 0.8693251 | 0.22352797 |
| Infected treated | PCR.lung | 0.9047046 | 0.40248488 |

Fisher’s F-test

| **Comparison** | **F** | **p-value** |
| --- | --- | --- |
| PCR_brain | 5.677e+10 | < 2.2e-16 |
| PCR_lung | 27426 | 9.816e-11 |

**Supplementary Table S6.** Ronapreve (RON), i.e. Casirivimab (CAS) and Imdevimab (IMD) concentration in the serum and tissues of SARS-CoV-2 Delta-infected and mock-infected K18-hACE2 mice that had received one intraperitoneal injection of RON (400 µg) and were euthanised 6 days later. In infected, untreated mice, the mAb concentrations were below the lower limits of quantification in serum and tissues.

| **Animal No** | **Infection** | **RON concentration (Ratio to SARS-CoV-2 Delta EC_90_)** | | | | | | |
| --- | --- | --- | --- | --- | --- | --- | --- | --- |
|  |  | **Serum**  **(ng/mL)** | **Brain**  **(ng/g)** | **Lung**  **(ng/g)** | **Nasal turbinate (ng/g)** | **Spleen**  **(ng/g)** | **Liver**  **(ng/g)** | **Kidney**  **(ng/g)** |
| 2.1 | Delta | CAS: 39,160 (946)  IMD: 29,933 (708) | CAS: 120 (2.90)  IMD: 260 (6.15) | CAS: 3,160 (76.3)  IMD: 2,950 (69.7) | CAS: 180 (4.35)  IMD: 310 (7.33) | CAS: 3,440 (83.1)  IMD: 3,950 (93.4) | CAS: 1,120 (27.1)  IMD: 1,630 (38.5) | CAS: 1,050 (25.4)  IMD: 2,230 (52.7) |
| 2.2 | Delta | CAS: 37,011 (894)  IMD: 24,697 (584) | CAS: 190 (4.59)  IMD: 380 (8.98) | CAS: 3,400 (82.1)  IMD: 3,030 (71.6) | CAS: 520 (12.6)  IMD: 670 (15.8) | CAS: 1,910 (46.1)  IMD: 3,150 (74.5) | CAS: 710 (17.2)  IMD: 1,760 (41.6) | CAS: 1,760 (42.5)  IMD: 2,560 (60.5) |
| 2.3 | Delta | CAS: 37,725 (911)  IMD: 30,201 (714) | CAS: 120 (2.90)  IMD: 300 (7.09) | CAS: 2,980 (72.0)  IMD: 2,560 (60.5) | CAS: 170 (4.11)  IMD: 310 (7.33) | CAS: 4,330 (104)  IMD: 3,820 (90.3) | CAS: 1,470 (35.5)  IMD: 2,500 (59.1) | CAS: 820 (19.8)  IMD: 2,160 (51.1) |
| 2.4 | Delta | CAS: 37,725 (911)  IMD: 35,144 (831) | CAS: 200 (4.83)  IMD: 340 (8.04) | CAS: 4,990 (120)  IMD: 3,260 (77.1) | CAS: 370 (8.94)  IMD: 500 (11.8) | CAS: 2,010 (48.6)  IMD: 2,490 (58.9) | CAS: 1,260 (30.4)  IMD: 2,060 (48.7) | CAS: 1,100 (26.6)  IMD: 2,070 (48.9) |
| 2.5 | Delta | CAS: 36,642 (885)  IMD: 23,754 (562) | CAS: 130 (3.14)  IMD: 300 (7.09) | CAS: 4,300 (104)  IMD: 3,200 (75.7) | CAS: 360 (8.70)  IMD: 500 (11.8) | CAS: 1,640 (39.6)  IMD: 2,550 (60.3) | CAS: 450 (10.9)  IMD: 2,040 (48.2) | CAS: 960 (23.2)  IMD: 2,620 (61.9) |
| 2.6 | Delta | CAS: 36,343 (878)  IMD: 32,633 (771) | CAS: 180 (4.35)  IMD: 340 (8.04) | CAS: 2,410 (58.2)  IMD: 2,250 (53.2) | CAS: 350 (8.45)  IMD: 470 (11.1) | CAS: 4,070 (98.3)  IMD: 3,410 (80.6) | CAS: 1,730 (41.8)  IMD: 2,650 (62.7) | CAS: 1,900 (45.9)  IMD: 3,100 (73.3) |
| 3.1 | Mock | CAS: 38,734 (936)  IMD: 32,119 (759) | CAS: 180 (4.35)  IMD: 350 (8.27) | CAS: 2,880 (69.6)  IMD: 1,940 (45.9) | CAS: 490 (11.8)  IMD: 510 (12.1) | CAS: 2,960 (71.5)  IMD: 3,070 (72.6) | CAS: 1,110 (26.8)  IMD: 1,770 (41.8) | CAS: 1,540 (37.2)  IMD: 2,360 (55.8) |
| 3.2 | Mock | CAS: 39,374 (951)  IMD: 38,206 (903) | CAS: 230 (5.56)  IMD: 370 (8.75) | CAS: 3,570 (86.2)  IMD: 1,830 (43.3) | CAS: 410 (9.90)  IMD: 530 (12.5) | CAS: 3,870 (93.5)  IMD: 2,080 (49.2) | CAS: 1,940 (46.9)  IMD: 2,680 (63.4) | CAS: 2,160 (52.2)  IMD: 2,390 (56.5) |
| 3.3 | Mock | CAS: 38,348 (926)  IMD: 34,146 (807) | CAS: 210 (5.07)  IMD: 330 (7.80) | CAS: 4,920 (118)  IMD: 3,680 (87.0) | CAS: 620 (14.9)  IMD: 650 (15.4) | CAS: 3,410 (82.4)  IMD: 2,720 (64.3) | CAS: 1,700 (41.1)  IMD: 2,280 (53.9) | CAS: 2,300 (55.6)  IMD: 2,650 (62.7) |
| 3.4 | Mock | CAS: 38,806 (937)  IMD: 29,847 (706) | CAS: 190 (4.59)  IMD: 320 (7.57) | CAS: 3,540 (85.5)  IMD: 2,990 (70.7) | CAS: 440 (10.6)  IMD: 530 (12.5) | CAS: 2,210 (53.4)  IMD: 4,460 (105) | CAS: 1,670 (40.3)  IMD: 2,440 (57.7) | CAS: 2,920 (70.5)  IMD: 3,460 (81.8) |

**Supplementary Table S7.** Relevant histological changes and SARS-CoV-2 nucleoprotein expression in K18-hACE-2 mice after intranasal infection with 10^3^ PFU SARS-CoV-2 Delta and euthanised at 7 days post infection (dpi). Animals in cohort 1 (n=5) were not treated. Animals in cohort 2 (n=6) were treated with Ronapreve once, via intraperitoneal injection, at 1 dpi (24 h after intranasal challenge).

| **Animal No** | **Treatment** | **Histological changes and viral antigen expression (lung, brain)** | **Virology (PCR)^1^** |
| --- | --- | --- | --- |
| 1.1 | Untreated | **Lung (HE)**: a few small focal areas with activated type II pn, a few macrophages, some LC, rare NL and scattered deg cells, some desquamation; several vessels with mild to mod pv leukocyte infiltration  **vAg**: variably sized patches of alveoli with pos AEC | 4.49E+07 |
|  |  | **Brain (HE)**: mild mononuclear pv infiltration (brainstem, medulla)  **vAg**: widespread patchy neuronal infection (brainstem, hippocampus, medulla, one patch in cerebellar cortex | 2.17E+10 |
| 1.2 | Untreated | **Lung (HE)**: a few small focal areas with activated type II pn, a few macrophages, some LC, rare NL and scattered deg cells, some desquamation; several vessels with mild pv leukocyte infiltration  **vAg**: variably sized patches of alveoli with pos AEC | 9.56E+07 |
|  |  | **Brain (HE)**: mild perivascular mononuclear infiltration (brainstem to medulla)  **vAg**: widespread neuronal infection (brainstem, cortex, hippocampus, medulla, individual cells in cerebellar cortex) | 2.36E+10 |
| 1.3 | Untreated | **Lung (HE)**: multiple, partly large focal, mod consolidated areas with activated type II pn, a few macrophages, several LC, rare NL and scattered deg cells, some desquamation; many vessels with mod pv leukocyte infiltration and evidence of leukocyte recruitment  **vAg**: disseminated small patches of alveoli with pos AEC | 1.63E+07 |
|  |  | **Brain (HE)**: NHA  **vAg**: neg | 3.47E+06 |
| 1.4 | Untreated | **Lung (HE)**: multiple, partly large focal, mod consolidated areas with activated type II pn, a few macrophages, several LC, rare NL and scattered deg cells, some desquamation; many vessels with mod pv leukocyte infiltration and evidence of leukocyte recruitment  **vAg**: numerous mainly small patches of alveoli with pos AEC | 3.21E+07 |
|  |  | **Brain (HE)**: NHA  **vAg**: neg | 5.83E+07 |
| 1.5 | Untreated | **Lung (HE)**: multiple, partly consolidated areas with activated type II pn, a few macrophages, several LC, rare NL and scattered deg cells, some desquamation; several vessels with mild pv leukocyte infiltration and evidence of leukocyte recruitment [PHOTOS?]  **vAg**: numerous mainly small patches of alveoli with pos AEC | 7.15E+07 |
|  |  | **Brain (HE)**: mild mononuclear pv infiltration (brainstem, medulla)  **vAg**: clusters of positive neurons (brainstem, medulla)  NB: Cortex or hippocampus not on section | 3.69E+09 |
| 2.1 | Ronapreve | **Lung (HE)**: mf small parenchymal areas of leukocyte aggregates (LC, macrophages, rare NL); several vessels with mild pv mononuclear infiltration  **vAg**: neg | 4.39E+05 |
|  |  | **Brain (HE)**: NHA  **vAg**: neg | 1.10E+05 |
| 2.2 | Ronapreve | **Lung (HE)**: mf small parenchymal areas of leukocyte aggregates (LC, macrophages, rare NL); several vessels with mild to mod pv mononuclear infiltration  **vAg**: neg | 6.33E+05 |
|  |  | **Brain (HE)**: NHA  **vAg**: neg | 1.45E+05 |
| 2.3 | Ronapreve | **Lung (HE)**: scattered small parenchymal leukocyte aggregates (LC, macrophages, rare NL)/granulomatous infiltrates; one larger similar area and mod mf pv mononuclear, LC-dom infiltration  **vAg**: neg | 4.51E+05 |
|  |  | **Brain (HE)**: NHA  **vAg**: neg | 5.44E+04 |
| 2.4 | Ronapreve | **Lung (HE)**: scattered small parenchymal leukocyte aggregates (LC, macrophages, rare NL)/granulomatous infiltrates; mild to mod mf pv mononuclear, LC-dom infiltration  **vAg**: neg | 5.74E+05 |
|  |  | **Brain (HE)**: NHA  **vAg**: neg | 4.69E+04 |
| 2.5 | Ronapreve | **Lung (HE)**: several small parenchymal leukocyte aggregates (LC, macrophages, rare NL)/granulomatous infiltrates; mild to mod mf pv mononuclear, LC-dom infiltration  **vAg**: neg | 3.81E+05 |
|  |  | **Brain (HE)**: NHA  **vAg**: neg | 2.28E+04 |
| 2.6 | Ronapreve | **Lung (HE)**: several small parenchymal leukocyte aggregates (LC, macrophages, rare NL)/granulomatous infiltrates; mod to marked mf pv mononuclear, LC-dom infiltration  **vAg**: neg | 8.72E+04 |
|  |  | **Brain (HE)**: NHA  **vAg**: neg | 2.25E+04 |

**Legend**: AEC – alveolar epithelial cells; AM – alveolar macrophages; deg – degenerate; dom – dominated; HE – histological features assessed in a hematoxylin-eosin stained section; in – intranasal; LC – lymphocyte; mf – multifocal; mod – moderate(ly); neg – negative; NHA – no histological abnormality; NL – neutrophils; LC – lymphocytes; LC-dom – lymphocyte dominated; occ – occasional; pn – pneumocytes; pos – positive; pv – perivascular; vAg – viral antigen

^1^ PCR: Copies of viral N1 RNA/µg of RNA relative to 18S, for each animal determined in lungs and brain (frontal cortex)

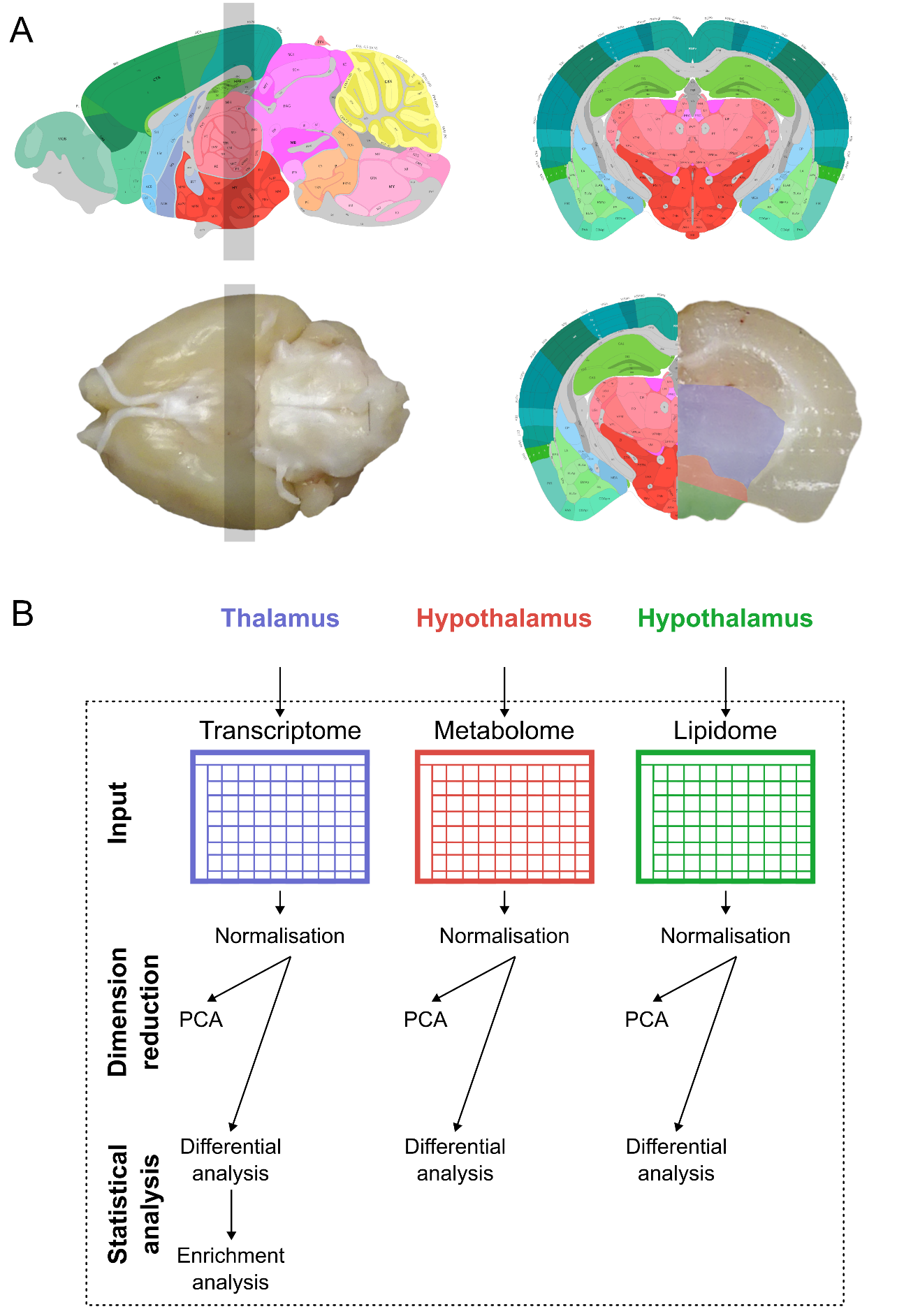

**Supplemental Figure S1**. Overview of the multiomics pipeline. **A)** Illustration of the tissue slice prepared by coronal sections (grey rectangle) to provide the samples for transmission electron microscopy and for the bulk transcriptomic, proteomic, metabolomic, and lipidomic analyses. Pictures of an example brain are mapped to Allen Mouse Brain Atlas (Allen Reference Atlas – Adult Mouse Coronal Sections (image 75), Mouse, P56 Sagittal (image 21) - <https://atlas.brain-map.org/>) (*33–36*). **B)** Overview of the workflow used for the single -omics analyses. PCA: principal component analysis.

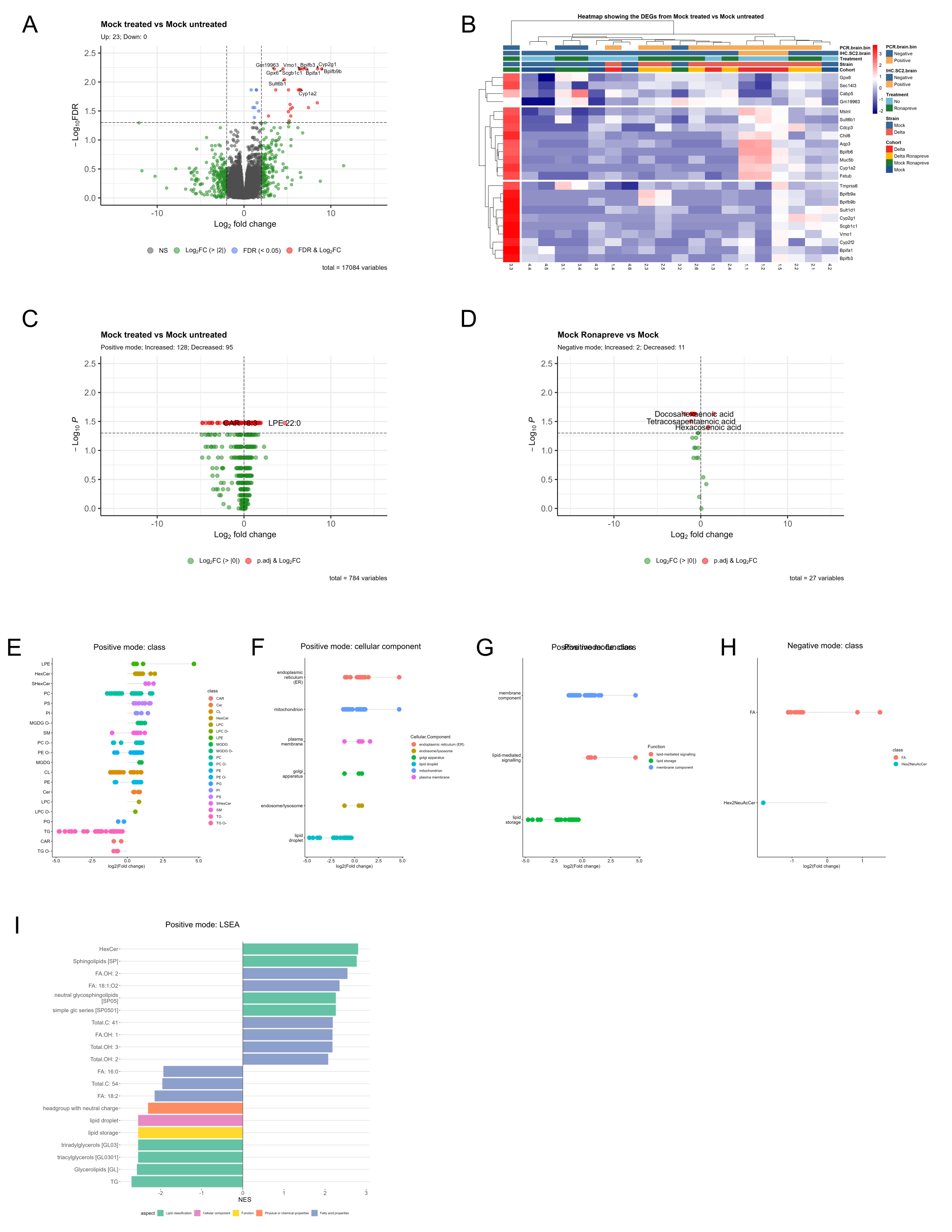

**Supplemental Figure S2. Results of the transcriptomic and lipidomic analyses, for the mock infected treated (mock treated) – mock infected untreated (mock untreated) comparison pair, when animal 3.3 is included**. **A,B)** Transcriptomic (thalamus). A) Volcano plot showing 23 significantly up-regulated genes. B) Heatmap of the significantly up-regulated genes, showing that animal 3.3 is a clear outlier in the mock infected treated cohort. **C-I)** Lipidomic (hypothalamus), positive and negative modes. C,D) Volcano plots showing the significantly different lipids. E-H) Lipid characteristics of the significantly different lipids; analysis was done in R using LipidSigR. I) Lipid Set Enrichment Analysis (LSEA) on the significantly different lipids from the positive mode; analysis was done in R using LipidSigR.

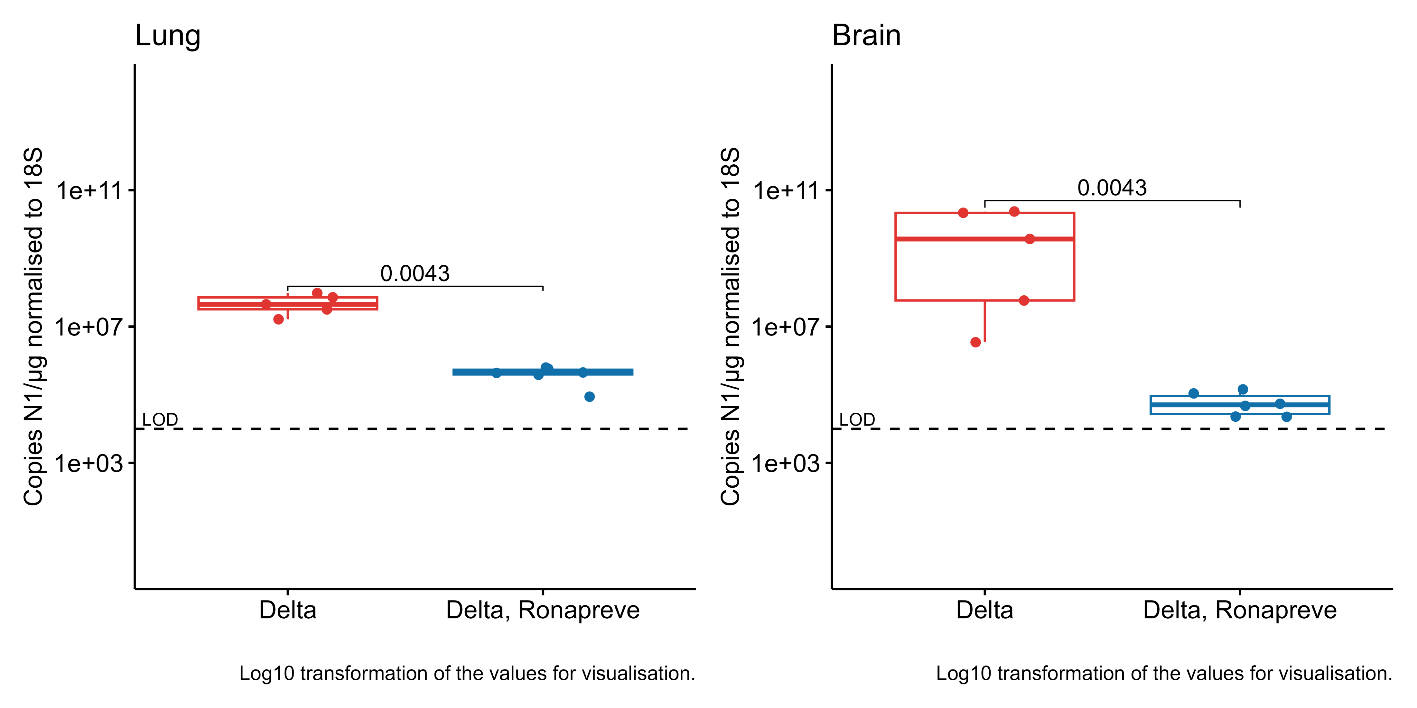

**Supplemental Figure S3**. Viral RNA loads in lung and brain of K18-hACE2 mice after intranasal challenge with SARS-CoV-2 Delta at 10^3^ PFU/mouse, examined at 7 dpi. SARS-CoV-2 N1 RNA was detected by RT-qPCR in lungs (A) and brains (B). With post-challenge treatment with Ronapreve (400 µg, ip) at 24 hpi, viral RNA levels significantly lower in both lungs and brain. LOD: limit of detection. The statistical comparisons were performed using a Mann-Whitney U test. The y axis was log10 transformed for the mean of visualisation only.

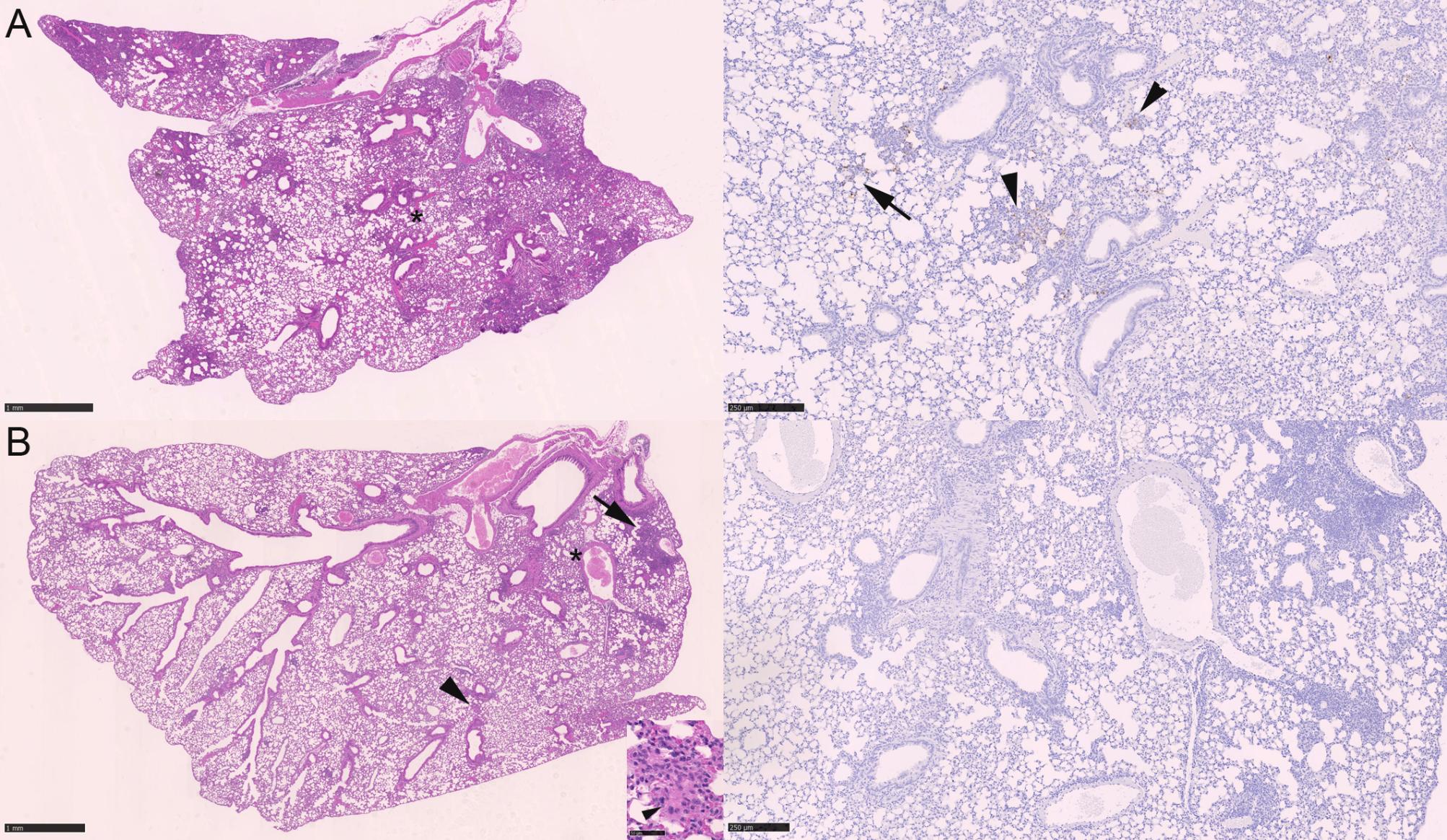

**Supplemental Figure S4**. Lungs, K18-hACE2 mice after intranasal challenge with SARS-CoV-2 Delta at 10^3^ PFU/mouse and treated with Ronapreve (400 µg, ip) at 24 hpi, examined at 7 dpi. A) Untreated animal. The overview (HE stain, left) shows multiple, moderately consolidated areas and moderate perivascular leukocyte infiltration and evidence of leukocyte recruitment. Staining for viral nucleoprotein (right; area indicated by asterisk in HE stain) highlights disseminated small patches of alveoli with positive alveolar epithelial cells, either unaltered (arrow) or within areas consolidated by leukocyte infiltrate (arrowheads). B) Ronapreve treated animal. The overview (HE stain, left) shows scattered, mainly small parenchymal leukocyte aggregates and granulomatous infiltrates (arrowhead, highlighted also in inset) and multifocal perivascular mononuclear infiltration (arrow). There is no evidence of viral nucleoprotein expression (right; area indicated by asterisk in HE stain). HE stain (left column); immunohistology for SARS-CoV-2 nucleoprotein, haematoxylin counterstain (right column). Bars: 1 mm (HE stain); 250 µm (NP stain); 50 µm (inset).

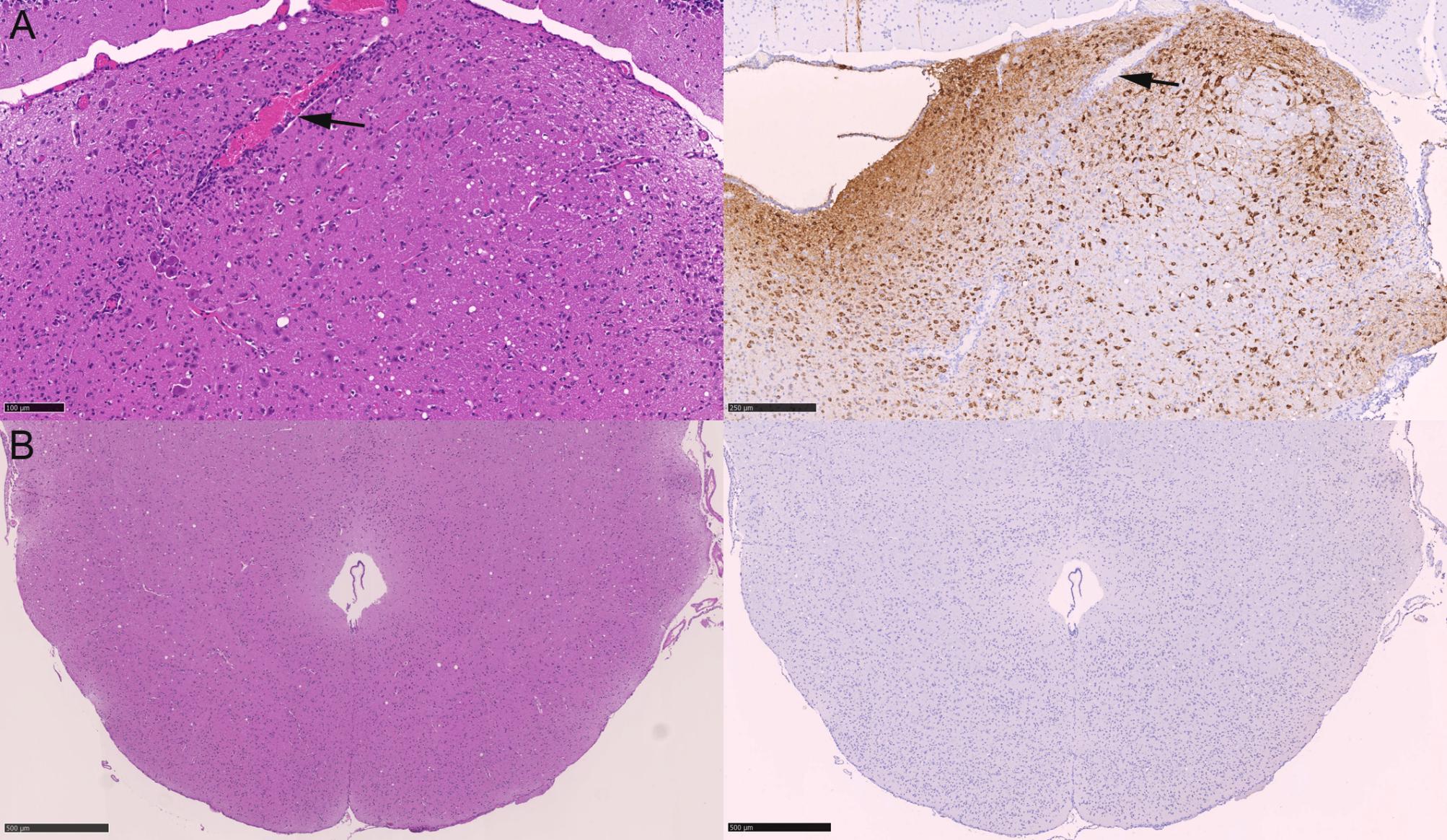

**Supplemental Figure S5**. Brainstem, K18-hACE2 mice after intranasal challenge with SARS-CoV-2 Delta at 10^3^ PFU/mouse and treated with Ronapreve (400 µg, ip) at 24 hpi, examined at 7 dpi. A) Untreated animal. The HE stain (left) shows a larger vessel (arrow) and some small vessels with a mild perivascular leukocyte infiltrate, consistent with a mild nonsuppurative encephalitis. Staining for viral nucleoprotein (right) highlights widespread neuronal infection. The arrow highlights the vessel shown in the HE stain. B) Ronapreve treated animal. The brainstem does not exhibit any histological changes (HE stain, left), and there is no evidence of viral nucleoprotein expression (right). HE stain (left column); immunohistology for SARS-CoV-2 nucleoprotein, haematoxylin counterstain (right column). Bars: 250 µm (A); 500 µm (B).

11.	Hervé Pagès, Marc Carlson, Seth Falcon, Nianhua Li, *AnnotationDbi* (Bioconductor, 2025).

12.	Marc Carlson, *org.Mm.eg.db* (Bioconductor, 2026).

13.	H. Wickham, *ggplot2*, *Elegant Graphics for Data Analysis* (Springer International Publishing; Imprint: Springer, Cham, ed. 2, 2016).

14.	H. Wickham, R. François, L. Henry, K. Müller, D. Vaughan, *dplyr: A Grammar of Data Manipulation* (2025).

15.	T. L. Pedersen, *patchwork: The Composer of Plots* (2025).

16.	H. Wickham, *stringr: Simple, Consistent Wrappers for Common String Operations* (2025).

17.	A. Lex, N. Gehlenborg, H. Strobelt, R. Vuillemot, H. Pfister, UpSet: Visualization of Intersecting Sets. *IEEE transactions on visualization and computer graphics*. **20**, 1983–1992 (2014), doi:10.1109/TVCG.2014.2346248.

18.	K. Müller, H. Wickham, *tibble: Simple Data Frames* (2026).

19.	Kolde Raivo, *pheatmap: Pretty heatmaps* (2025).

20.	Guangchuang Yu, *enrichplot* (Bioconductor, 2025).

21.	X. Shen *et al.,* TidyMass an object-oriented reproducible analysis framework for LC-MS data. *Nature communications*. **13**, 4365 (2022), doi:10.1038/s41467-022-32155-w.

22.	A. Kassambara, *rstatix: Pipe-Friendly Framework for Basic Statistical Tests* (2025).

23.	H. Wickham, D. Vaughan, M. Girlich, *tidyr: Tidy Messy Data* (2025).

30.	Martin Morgan, Valerie Obenchain, Jim Hester, Hervé Pagès, *SummarizedExperiment* (Bioconductor, 2025).

31.	R. M. Hope, *Rmisc: Ryan Miscellaneous* (2022).

32.	A. Kassambara, *ggpubr: 'ggplot2' Based Publication Ready Plots* (2025).

33.	T. L. Daigle *et al.,* A Suite of Transgenic Driver and Reporter Mouse Lines with Enhanced Brain-Cell-Type Targeting and Functionality. *Cell*. **174**, 465-480.e22 (2018), doi:10.1016/j.cell.2018.06.035.

34.	E. S. Lein *et al.,* Genome-wide atlas of gene expression in the adult mouse brain. *Nature*. **445**, 168–176 (2007), doi:10.1038/nature05453.

35.	J. A. Harris *et al.,* Hierarchical organization of cortical and thalamic connectivity. *Nature*. **575**, 195–202 (2019), doi:10.1038/s41586-019-1716-z.

36.	S. W. Oh *et al.,* A mesoscale connectome of the mouse brain. *Nature*. **508**, 207–214 (2014), doi:10.1038/nature13186.
